# Disruption of Circadian Clock Induces Abnormal Mammary Morphology and Aggressive Basal Tumorigenesis by Enhancing LILRB4 Signaling

**DOI:** 10.1101/2024.03.19.585534

**Authors:** Olajumoke Ogunlusi, Mrinmoy Sarkar, Arhit Chakrabarti, Devon J Boland, Tristan Nguyen, James Sampson, Christian Nguyen, Danielle Fails, Yava Jones-Hall, Loning Fu, Bani Mallick, Alex Keene, Jeff Jones, Tapasree Roy Sarkar

## Abstract

Epidemiological studies have shown that circadian rhythm disruption (CRD) is associated with the risk of breast cancer. However, the role of CRD in mammary gland morphology and aggressive basal mammary tumorigenesis and the molecular mechanisms underlying CRD and cancer risk remain unknown. To investigate the effect of CRD on aggressive tumorigenesis, a genetically engineered mouse model that recapitulates the human basal type of breast cancer was used for this study. The effect of CRD on mammary gland morphology was investigated using wild-type mice model. The impact of CRD on the tumor microenvironment was investigated using the tumors from LD12:12 and CRD mice via scRNA seq. ScRNA seq was substantiated by multiplexing immunostaining, flow cytometry, and realtime PCR. The effect of LILRB4 immunotherapy on CRD-induced tumorigenesis was also investigated. Here we identified the impact of CRD on basal tumorigenesis and mammary gland morphology and identified the role of LILRB4 on CRD-induced lung metastasis. We found that chronic CRD disrupted mouse mammary gland morphology and increased tumor burden, and lung metastasis and induced an immunosuppressive tumor microenvironment by enhancing LILRB4a expression. Moreover, CRD increased the M2-macrophage and regulatory T-cell populations but decreased the M1-macrophage, and dendritic cell populations. Furthermore, targeted immunotherapy against LILRB4 reduced CRD-induced immunosuppressive microenvironment and lung metastasis. These findings identify and implicate LILRB4a as a link between CRD and aggressive mammary tumorigenesis. This study also establishes the potential role of the targeted LILRB4a immunotherapy as an inhibitor of CRD-induced lung metastasis.

## Introduction

The circadian clock regulates the expression of several genes that impact physiology, metabolic processes, and health outcomes (Panda, Antoch et al. 2002, Storch, Lipan et al. 2002, Kelleher, Rao et al. 2014). Epidemiological studies have reported that circadian rhythm disruption (CRD), such as that which occurs during shift work or travel across time zones, affects human health and increases the risk of developing cancer, metabolic disorders, and cardiovascular disease (Hemmer, Mareschal et al. 2021). According to the National Health Interview Survey, 12–35% of the US population works irregular schedules, including night and rotating shifts (CDC 2015). CRD increases the risk of different types of cancer, including lung (Papagiannakopoulos, Bauer et al. 2016, Pariollaud, Ibrahim et al. 2022), colon (Chun, Fortin et al. 2022), and breast (Van Dycke, Rodenburg et al. 2015, Hadadi 2020) cancers.

Shiftwork was found to increase the incidence of breast cancer in nurses by approximately 50%, suggesting a critical role for the circadian clock in breast cancer pathogenesis (Schernhammer, Laden et al. 2001). Circadian clocks regulate the rhythmic expression of numerous genes in breast tissues (Blakeman, Williams et al. 2016). Triple-negative breast cancer (TNBC) or basal phenotype (Alluri and Newman 2014) encompasses a breast tumor subtype that is clinically negative for the expression of the estrogen (ER) and progesterone (PR) receptors and lacks overexpression of the Human Epidermal Growth Factor Receptor 2 (HER2) protein, with unique prognostic and therapeutic implications (Anders and Carey 2008). TNBC is responsible for more than 15–20% of all breast cancers, with a mortality rate of 40% (Almansour 2022). TNBC is relatively aggressive and shows more resistance to chemotherapy than other subtypes of breast cancer (Almansour 2022). However, the effect of CRD on aggressive TNBC is poorly understood, and the precise mechanism underlying CRD-induced tumorigenesis is not yet known.

The mammalian circadian machinery consists of an autoregulatory transcription-translation feedback loop, where the “positive elements” circadian locomotor output cycle kaput (CLOCK) and brain and muscle aryl hydrocarbon receptor nuclear translocator (ARNT)-like protein 1 (BMAL1) heterodimerize through their PAS domains and activate the transcription of the “negative elements,” i.e., the period genes (*Per1* and *Per2)* and the cryptochrome genes (*Cry1* and *Cry2)*. The PER/CRY heterodimers inhibit the transcription of their gene(s) by blocking CLOCK/BMAL1-dependent transactivation (Kelleher, Rao et al. 2014). This feedback loop regulates the transcription of an estimated 30% of the genome (Vitaterna, Shimomura et al. 2019). The expression of core circadian clock genes is frequently dysregulated in human tumors, indicating the tumor-suppressive role of the molecular clock (Battaglin, Chan et al. 2021, Stephenson, Usselmann et al. 2021, Chun, Fortin et al. 2022). Epidemiological studies have also emphasized that the risk of developing cancer increases with increasing years of shift work (Yuan, Zhu et al. 2018), indicating that longitudinal experiments are better suited for characterizing how CRD may affect tumorigenesis.

Here, we investigated how the loss of circadian function impacts mammary glands, tumorigenesis, and the tumor microenvironment (TME). We used the jet lag schedule, which mimics the circadian disturbance humans encounter during rotating shift work or frequent jet lag, and the genetically engineered mouse model (GEMM) of TNBC. We specifically examined the role of leukocyte immunoglobulin-like receptor 4a (LILRB4a or LILRB4 or gp49B), which is known to suppress immunity in acute myeloid leukemia (AML) and solid tumors (Deng, Gui et al. 2018, Sharma 2021). Additionally, we investigated whether the elevation of LILRB4 expression could be a key molecular event in CRD-induced mammary tumorigenesis and immunosuppressive TME. Overall, this study shows how circadian desynchronization affects mammary gland morphology, aggressive mammary tumorigenesis, and identifies a compelling target for immunotherapy of CRD-induced tumorigenesis.

## Methods

### A. Animal husbandry

The mice were housed under 12 h light and 12 h dark (LD 12:12) and circadian rhythm disruption (CRD) conditions at room temperature (25°C), and food and water were provided *ad libitum*. The genetically engineered mice model (GEMM), i.e., FVB-Tg(C3-1-Tag)cJeg/JegJ mice (Jax stock #013591), were caged until they were 27 weeks old. Normal FVB mice (Jax stock #001800) and BALB/cJ mice (Jax stock #000651) were also used in this study. All animal care and treatments were per the Institutional Animal Care and Use Committee of Texas A&M University under protocol # 2022-0094.

Mice were grouped randomly for exposure to standard light conditions (LD 12:12) or CRD light conditions (consisting of an 8-hour light phase advance repeated every two days) (Casiraghi, Oda et al. 2012, Papagiannakopoulos, Bauer et al. 2016, Pariollaud, Ibrahim et al. 2022). Zeitgeber time (ZT) 0 corresponded to the onset of light, while ZT12 corresponded to the onset of dark. The LD 12:12 and CRD animal groups were provided access to running wheels (Columbus Instruments) for several weeks. ActogramJ (ImageJ) was used to assess voluntary running activities (Schmid, Helfrich-Förster et al. 2011).

### B. 4T1 cell mouse TNBC model

The 4T1 cells were maintained RPMI-1640 (ATCC, 30-2001) supplemented with in 10% FBS and 1% penicillin and streptomycin in 5% CO_2_. Briefly, 10,000 cells suspended in sterile PBS were injected subcutaneously into the 4^th^ mammary gland fat pad. The mice were housed in LD or CRD conditions for 3-4 weeks before they were sacrificed to harvest the tumors. To investigate the effect of immunotherapy on tumor progression and metastasis, BALB/cJ mice were housed under LD 12:12 conditions for 4-5 days following injection of 4T1 cells until palpable tumors developed. They were segregated and assigned randomly either in LD or CRD conditions. On days 6, 9, and 12 (after the palpable tumors developed), an anti-LILRB4 antibody (ABclonal, #A7073) (polyclonal, 50μg) was injected intratumorally. The animals were checked regularly to observe their body weight and tumor sizes.

### C. Histology and tumor burden analysis

The tumors and the mammary glands were harvested using aseptic techniques. Tumor volumes were calculated for each breast tumor using caliper measurements after every 3 days. *Tumor volume* = (*length* × *width*^2)/2 was used to measure the volume of the tumors (Kersemans, Cornelissen et al. 2013), whereas the equation *Tumor burden* (*in percentage*) = (*Tumor weight* /*mice body weight*) × 100 % was considered to determine the tumor burden (Hadadi 2020). The tumor, or the lung samples were fixed in 10% neutral buffered formalin (NBF) for 48 hours before being transferred to the Histology core facility at Texas A&M University. The tissue sections were stained with hematoxylin and eosin (H&E) for pathological examinations.

### D. Immunohistochemistry

The formalin-fixed paraffin-embedded (FFPE) mouse mammary gland tissues were deparaffinized by incubating at 65°C for 2 hours, followed by dipping in xylene for 5 minutes each time for two changes. The endogenous peroxidase activity was inhibited by treating the tissue with hydrogen peroxide. Antigen retrieval was carried out by incubating the slides in pre-heated citrate antigen retrieval buffer (10 mM citrate, 0.05% Tween 20, pH 6.0) at 95°C for 20-25 minutes in a water bath. After blocking, the tissues were incubated overnight at 4°C with anti-ki67 polyclonal antibody (ABclonal, A16919), dissolved in the kit blocking solution. Following incubation with the secondary antibody availed with the IHC kit, the expression was visualized with DAB peroxidase substrate (Abcam, ab64264). The slides were counterstained with Mayer’s Hematoxylin Solution (VitroVivo Biotech, VB-3000-1) before being mounted with Optic Mount I™ xylene mounting medium (Mercedes Scientific) and imaged under the Echo Revolve R4 fluorescent microscope (Echo).

### E. Whole mount staining of mouse mammary gland

The mammary gland tissues were spread on glass slides immediately after harvesting. The mammary glands were then fixed using Carnoy’s fixative solution (VitroVivo Biotech, VB-3001-1) for three hours, followed by washing in 70% ethanol for 30 minutes. The tissues were incubated overnight at room temperature (RT) with Carmine Alum Staining Solution (VitroVivo Biotech, VB-3001-2). Stained tissues were dehydrated with gradients of ethanol concentrations and finally with three changes of xylene bath of 10 minutes each. The slides were then mounted with Optic Mount I™ xylene mounting medium (Mercedes Scientific) and observed under the microscope.

### F. Single-cell RNA sequencing (scRNA Seq)

After brief washing in chilled sterile PBS, the tumors were kept in DMEM (ATCC, 30-2002), supplemented with 10% FBS (Sigma-Aldrich, F2442) and 1% antibiotic solution (Sigma Aldrich, P4333) on ice until minced and subsequently blended with gentleMACS Dissociator (Miltenyi Biotec, 130-093-235). MACS Miltenyi Tumor Dissociation Kit for mice (Miltenyi Biotech, 130-096-730) was used for further enzymatic digestion according to the manufacturer’s protocol. The suspension of cells was then transferred to the Texas A&M Institute for Genome Sciences and Society (TIGSS) for scRNA-seq. Single-cell sequencing libraries were prepared on the Chromium platform (10x Genomics) using the Single Cell 5’ v2 Full Kit (PN-1000263). Cell type assignment was performed manually based on unique marker genes identified in each cell cluster from the UMAP reduction. Markers were cross-referenced with published literature specific to mouse mammary gland tissues. Differentially expressed genes were identified using the MAST (Finak, McDavid et al. 2015) R package, and GO pathway enrichment analysis was performed with the clusterProfiler (Wu, Hu et al. 2021) R package. The CellChat R package analyzed cellular communication using default parameters. The most significant pathways were identified using the “rankNet” function and visualized with the “netVisual_diffInteraction” function.

### G. *In vitro* tumorosphere formation assay

The single cells (from tumors) were cultured in ultra-low attachment 96-well plates (ThermoFisher Scientific, 174927) in EpiCult™-B (mouse) medium supplemented with human recombinant EGF (hrEGF) (Stem Cell Technologies, 78006), human recombinant bFGF (hrbFGF) (Stem Cell Technologies, 78003), and heparin (Stem Cell Technologies, 07980). 500 cells per well from LD 12:12 and CRD group tumors were added and incubated with standard cell culture conditions for 5 days in the CO_2_ incubator before being observed under the microscope.

### H. Organoid assay

Singlet cells from tumors (LD and CRD) were resuspended in Geltrex™ (Gibco, ThermoFisher, A1413202) and plated on a 24-well plate. After the gel was solidified to form the matrix, EpiCult™-B (mouse) medium supplemented with hrEGF, hrbFGF, and heparin was added to assess the growth of the mouse mammary tumor cells from LD 12:12 and CRD conditions. After 7 days of incubation, the organoids formed were imaged under the microscope.

### I. Flow cytometry

The dissociated tumor cells were counted and resuspended in 1 ml of cell staining buffer (CSB) (BioLegend, 420201). Zombie NIR™ Fixable Viability Kit (BioLegend, 423105) was applied to cells on ice for 20 min. After spinning down at 400 x g for 4 minutes, cells were resuspended in 50 µl of the CSB and incubated with fluorochrome-conjugated antibodies (Alexa Fluor® 700 anti-mouse CD4, Pacific Blue™ anti-mouse CD8a, Alexa Fluor® 488 anti-mouse CD86, PE anti-mouse CD163, Alexa Fluor® 594 anti-mouse CD45, FITC anti-mouse Ly-6G/Ly-6C, PE anti-mouse/human CD11b, Alexa Fluor® 647 anti-mouse I-A/I-E, FITC anti-mouse CD11c) in the dark for 20 minutes, following a systematic approach via designing compatible multicolor flow cytometry panels. Cells were fixed and permeabilized using 2% paraformaldehyde and 0.1% Triton X-100 in PBS, respectively, and stained with anti-FoxP3 antibody (Alexa Fluor® 488 anti-mouse/rat/human FOXP3) for 20 minutes in the dark at 4°C, in case of tumor infiltrated lymphocyte-specific panel. Afterward, stained cells were rinsed with PBS and kept in PBS at 4°C in the dark until analyzed using Cytek Aurora spectral flow cytometer at TAMU Flow Cytometry Facility.

### J. Quantitative PCR for detecting mRNA levels

Total RNA was extracted from mammary glands or tumors using Quick-RNA™ MiniPrep (Zymo Research, R1054), and the purity was analyzed by DeNovix DS-11 Series nanodrop Spectrophotometer (DeNovix). RNA from each sample was reverse transcribed to cDNA by High-Capacity cDNA Reverse Transcription Kit (Applied Biosystems™, 4368814). Analysis for gene expression by quantitative real-time PCR was performed using SsoAdvanced Universal SYBR Green Supermix (BioRad, 1725271).

### K. Human CRD analysis for breast cancer patients

All statistical analyses were performed using the R software package (version 4.3.0.). scRNA-seq data corresponding to the triple-negative breast cancer (TNBC) patients from Wu *et al*., 2021 (Wu, Al-Eryani et al. 2021) were pre-processed using standard steps outlined in Hao *et al.,* (Hao, Hao et al. 2021). The uniform manifold approximation and projection (UMAP) was performed on the processed data. The UMAP plot with cells clustered according to the inferCNV cell type. The cells which had no infer-CNV call are denoted as “unknown.” To investigate the level of circadian rhythm disruption in each cell, we calculated the CRD scores for the 358 circadian-related genes (CRGs) following He *et al.,* 2022 (He, Fan et al. 2022) separately for malignant and non-malignant cells. A cutoff of 75% based on quartiles was used as the threshold for the CRD level to segregate the scores as high (denoted as CRD^hi^) and low (denoted as CRD^low^). The percentage of cells that show CRD^hi^ scores by cell type, along with the p-value of the corresponding Binomial proportion test, is used for this study. The mean normalized expression of the core circadian genes by the cell type (malignant and non-malignant) is presented in this study. Bulk RNA-sequencing data from the publicly available TCGA BRCA dataset was used further to explore the expression pattern of the core circadian genes. The patients were segregated according to the analyzed tissue type, i.e., “Tumor” and “Surrounding” tissue. The mean expression of the genes (+/- SEM) over patients by the type of tissue analyzed. The Wilcoxon signed-rank test was used to test for the difference in expression between the tumor and surrounding tissues for each gene. The genes showing a significant difference are labeled in “red”, while the insignificant genes are marked in “black”.

### L. Blood chemistry and cytokine array

Pre-mortem terminal blood was collected from the FVB-Tg(C3-1-Tag)cJeg/JegJ mice. The blood samples were transferred to the Texas A&M TVMDL to analyze the blood chemistry parameters. The cytokine array was performed using the Proteome Profiler Mouse XL Cytokine Array kit (R&D Systems, ARY028). The blood samples were kept undisturbed for 30 minutes at room temperature before being spun at 2000 x g for 10 minutes at 4°C. The serum was separated for the cytokine array profiling according to the manufacturer’s protocol and was scanned using ChemiDoc Imaging System (BioRad, 12003153).

### M. mIF Staining of Mouse Tissues

The panels were developed and optimized for use in mouse tissue. The following primary antibodies from Fortis Life Sciences, Cell Signaling Technology, and ABclonal were utilized for both immunohistochemical and immunofluorescence staining: rabbit anti-mouse ARG1 [BLR161J], rabbit anti-human/mouse LILRB4 [A7073], rabbit anti-human/mouse CD163 [BLR087G], rabbit anti-mouse CD8a [BLR173J], and rabbit anti-mouse FOXP3 [D6O8R], and rabbit anti-mouse CD4 [BLR167J]. Multiplex immunofluorescence staining was performed with the Akoya Opal™ Polaris 7-color IHC kit fluorophores (Akoya Biosciences [NEL861001KT]). FFPE tissue sections were baked for 30 minutes, deparaffinized in xylene, and rehydrated by serial passage through graded concentrations of ethanol. Endogenous peroxidase in tissues was blocked with 0.9% H_2_O_2_/methanol for 40 min. Multiple (7) sequential HIER treatments were performed for 20 min each at 92-96°C in Citrate pH 6.0 (first cycle) or Tris EDTA pH9 (remaining cycles) buffer. After each HIER cycle, sections were rinsed with DI water and cooled at RT for 20 minutes. Tissue sections were blocked with 20% normal goat serum for 20 minutes before incubation with primary antibody for 20 min. Then, sections were rinsed with TBST for 10 min and incubated with HRP-conjugated secondary (A120-501P) for 20 min, followed by another 10 min rinse in TBST. Incubation with an Opal fluorophore (Opal480, Opal520, Opal570, Opal620, Opal690) was then done for 10 min, followed by a 10 min rinse in DI water. Bound primary and secondary antibodies were then subject to the next HIER treatment (as aforementioned) for 20 min. After washing in DI water and cooling at RT for 20 min, the process of staining and antibody removal was repeated for the desired number of targets. Finally, the tissue specimens were stained with 4′,6-diamidino-2-phenylindole (DAPI) for 10 min and mounted in VECTASHIELD Vibrance Antifade Mounting Medium (ThermoFisher Scientific). Akoya Biosciences’ PhenoImager HT was used for multispectral imaging at 40× magnification.

### N. Statistical analysis

GraphPad Prism8 was used as the Statistical analysis tool except for the assessment of the rhythmicity that was carried out using the JTK cycle. The significance was assessed using either an Unpaired t-test or analysis of variance (ANOVA), with a false positive threshold of 0.05 considered acceptable (p<0.05). Unless otherwise mentioned, the data were presented as mean ± standard error of mean (SEM).

## Results

### Chronic CRD results in abnormal mammary gland morphology

The jet-lag paradigm mimics the CRD that humans face during shift work (Lee, Donehower et al. 2010) or travels across time zones. Mice were exposed to an 8 h light advance repeated every 2–3 days (Fig. 1A). Female Friend LeukemiaVirus B (FVB) mice (4 weeks old) were housed under normal light and dark (LD 12:12), or chronic CRD conditions for 8 weeks before the mammary glands were collected every 4 h for 24 h. We confirmed that this jet-lag protocol disrupted circadian activity in our mouse model by assessing the locomotor activity (Fig. 1B). CRD did not significantly impact the weight of mice (Fig. S1A). Real-time PCR analysis showed that CRD led to the loss of rhythmicity (*P*JTKcycle < 0.05) for most genes, including *Cry1*, *Cry2*, *Bmal1*, *Clock,* and *Per1* (Fig. 1C). *Per2* expression remained rhythmic in mice exposed to CRD (Fig. 1C), suggesting the desynchronization of the core clock components. To understand the effect of CRD on the mammary glands, we examined whole-mount preparations of inguinal mammary glands from LD and CRD mice at 12 weeks. Chronic CRD decreased branching (bifurcations) (Fig. 1D, upper panel, 1E, S1B) and the number of terminal end buds (TEBs) (Fig. 1D, lower panel, 1F, S1B) in the mammary glands. Compared with the mammary glands of LD mice (control), CRD mice showed a high frequency of ductal hyperplasia as found by using hematoxylin and eosin (H&E) (Fig. 1G) and by using Ki67 (Fig. 1H, S1C) staining of the mammary glands.

**Fig. 1.**
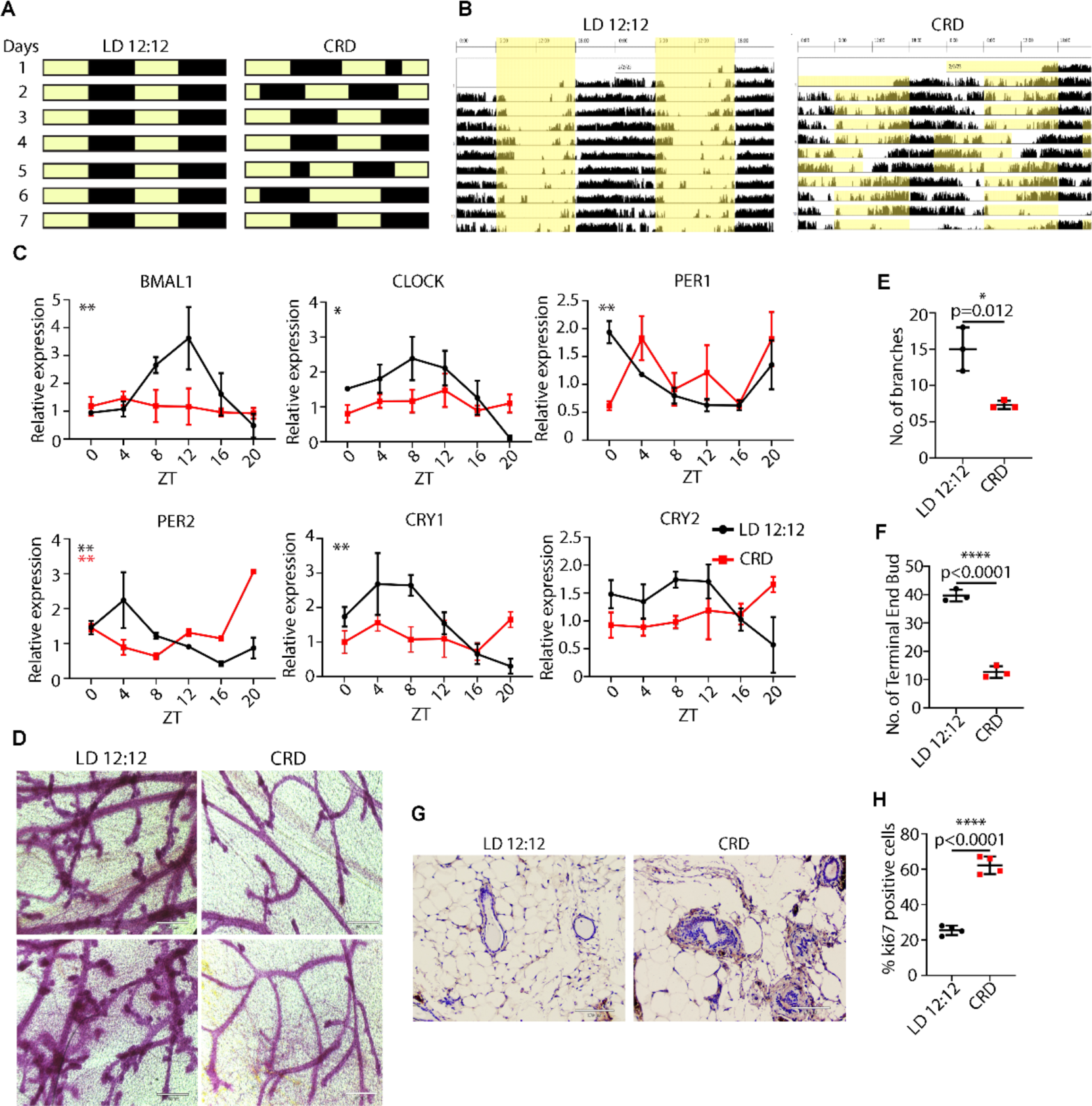
CRD severely impairs rhythmicity and disrupts mammary gland morphogenesis. **(A)** Schematic representation of the CRD protocol. **(B)** FVB female mice were housed under LD 12:12 or CRD for 8 weeks. Representative actograms of LD and CRD mice. LD: 12 h light and 12 h dark; CRD: circadian rhythm disruption using jet lag protocol, represented by a shortening of the dark period by 8 h every third day. **(C)** Mammary glands were collected at the indicated times. Gene expression normalized to GAPDH expression was measured using quantitative real-time PCR data represented as mean ± SEM for n=3 females per time point and light condition. Rhythmicity was determined using JTK_Cycle analyses; \**P*JTKcycle < 0.05, \*\**P*JTKcycle < 0.01, \*\*\**P*JTKcycle < 0.001, and \*\*\*\**P*JTKcycle < 0.0001. CRD disrupts branching morphogenesis in WT virgin FVB mice. **(D, E, F)** Representative images of branching and development of terminal ducts in LD and CRD animals are shown in LD and CRD-induced mammary glands **(D, E, F)**, scale bar: 350 μm, (\**P* < 0.05, error bars indicate SEM, n = 3). **(G)** The effect of CRD on mammary ductal hyperplasia is shown by hematoxylin and eosin (H&E) staining of the mammary gland (**G**) and by using Ki67 **(H)** of the mammary glands. Scale bar: 100 μm.

### Chronic CRD accelerates aggressive mammary tumorigenesis and lung metastasis

To investigate the effect of CRD on aggressive basal tumorigenesis, we used preclinical GEMM of human basal tumors (FVB-Tg(C3-1-Tag)cJeg/JegJ (Maroulakou, Anver et al. 1994, Usary 2017) that provides a model for examining how CRD impacts aggressive mammary tumorigenesis and metastasis. Female hemizygous mice develop mammary gland adenocarcinomas at approximately 24 weeks of age (Maroulakou, Anver et al. 1994, Usary 2017). To investigate the effect of CRD on tumorigenesis, mice were placed under LD or CRD conditions when they were 8 weeks old and maintained under CRD conditions through the end of the study at 27 weeks (Fig. 2A). The growth of primary tumors was measured every 3 days throughout this period. Because of the highly aggressive tumors in CRD mice, we euthanized the animals at 27 weeks. All mice were euthanized, and the tumor tissues were processed for other analyses. CRD did not significantly impact the weight of mice (Fig. S2A). Peripheral blood cell counts showed no significant differences between those mice (Fig. S2B, S2C). No significant changes were noted in the blood glucose, cholesterol, or amylase (Fig. S2D-F) levels in CRD mice, although the albumin-to-globulin ratio (AGR) was decreased (Fig. S2G). A high AGR shows a good prognosis for survival in different solid-tumor patients (He, Pan et al. 2017). Therefore, CRD negatively affects cancer prognosis by decreasing the AGR. scRNAseq (Fig. S3A) and real-time PCR (Fig. S3B) of tumor samples showed that CRD did not substantially alter the expression of Per genes (*Per1*, and *Per2*) (Fig. S3A, B) or other clock genes (*Clock*, *Cry2*) (Fig. S3A, B). *Cry1* expression significantly increased in CRD-induced tumors (Fig. S3B).

**Fig. 2.**
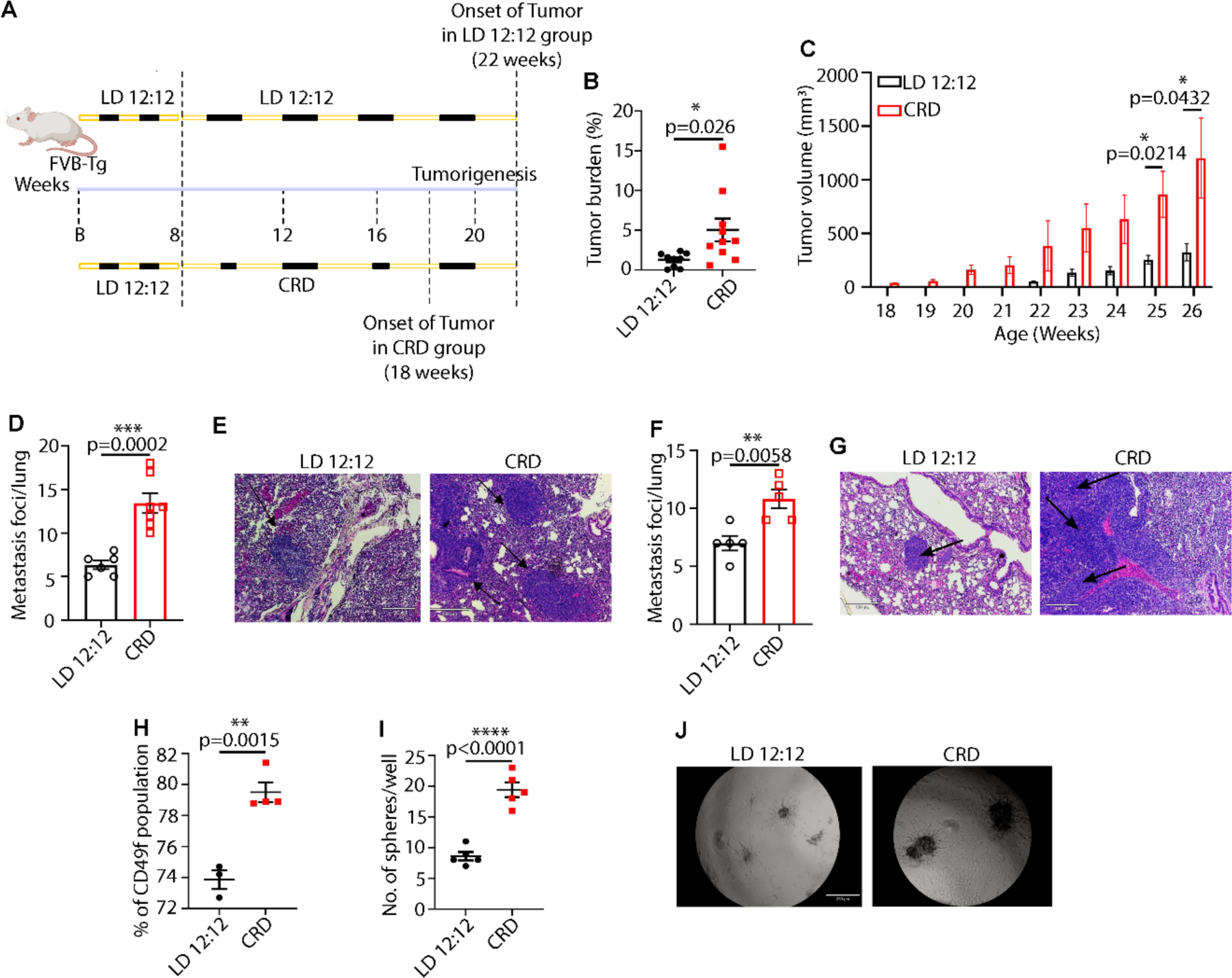
CRD increases aggressive mammary tumorigenesis and lung metastasis. Experimental timeline for evaluation of the effect of CRD on spontaneous TNBC in (FVB-Tg(C3-1-TAg)cJeg (C3-TAg) mice **(A)**. Dashed lines highlight the start and end points of the experiment. **(B)** Tumor burden (tumor to body weight ratio) as % in LD (n = 9) or CRD (n = 10) conditions. Column data represent the mean ± SEM values for individual animals. The effect of CRD on tumor initiation is shown in **(C)**. Number of metastatic foci in the lungs of LD (n=6) and CRD (n =7) mice at the time of their sacrifice is shown in **(D)**. P-value obtained from a binomial two-sided test. Indicated (n) represents the number of independent experiments as biological replicates. Metastatic foci in the lungs are shown using H&E staining, scale bar 150 μm **(E)**. CRD-induced lung metastasis in 4T1 tumor model in BALB/cJ mice as shown by the graphical representation **(F)** and H&E staining **(G),** scale bar 100 μm. **(H)** The percentage of CD49f-breast cancer stem cell (CSC) subpopulations in LD and CRD tumors is shown as a graph. Results represent the mean ± SEM of 3–4 independent experiments. P-value obtained from an unpaired two-sided *t*-test. Mammosphere formation efficiency (MFE%) of LD and CRD tumor cells (n =3) is shown in Figure. **(I)** with lines indicating the mean ± SEM. P-value obtained from an unpaired two-sided *t*-test. The formation of organoids (tumoroids) from LD and CRD tumors is shown in the **J**. Scale bar, 250 μm.

Animals placed in CRD conditions had a substantially increased mammary tumor burden (Fig. 2B) compared to the LD animals. CRD also accelerated tumor initiation at 18 weeks. In contrast, the average time of tumor initiation for LD mice was 22 weeks (Fig. 2C). However, no difference was observed in the spectrum of tumor grades assessed using histopathology (Elston and Ellis 2002) between the two groups, with most tumors being grade 2 (Fig. S4A, B). Several technical differences, such as in the mouse genetic background (MMTV:PyMT C57BL/6J versus FVB-Tg(C3-1-TAg)cJeg/JegJ) and conditions of the animal facilities (such as light exposure) may explain why Hadadi *et al*. (Hadadi 2020) observed an effect of CRD on tumor grade, but the same was not found in our study. The number of metastatic foci in the lungs was significantly higher in the CRD mice (Fig. 2D, E). To investigate the impact of CRD on tumor progression, 4T1 (mice triple-negative breast cancer cell line) cells were introduced into the female BALB/cJ mice via orthotopic transplantation into the 4th mammary glands. When mice developed palpable tumors (∼4-5 days after injection), they were placed into LD or CRD conditions for 3 weeks until tumors became aggressive. CRD led to a modest but non-significant increase in tumor burden (Fig. S4C). This data suggests that CRD does not affect mammary tumor progression. However, we did observe a significant increase in the formation of lung metastasis foci (Fig 2F, G). Flow cytometry was performed to investigate whether a higher metastatic potential was related to a higher cancer stem cell (CSC) population in CRD tumors. We observed a significant increase in the percentage of CSCs expressing CD49f markers (Fig. 2H, S4D) in CRD tumors compared with that in LD tumors. To investigate whether this enrichment was associated with an increase in the stemness potential of primary tumors, a mammosphere assay was performed using CRD and LD GEMM tumors. CRD tumor cells had a significantly higher (P<0.0001) mammosphere formation efficiency than LD tumor cells (Fig. 2I, S4E). Recent studies have used three-dimensional (3D) organoids as *in vitro* organs, with numerous applications ranging from disease modeling to drug screening (Meyer, Howden et al. 2011). Using small intestinal crypt-derived organoids, a recent study showed that genetic disruption of the circadian clock accelerates intestinal organoid proliferation (Chun, Fortin et al. 2022). We established 3D organoids (tumoroids) from LD and CRD tumors (Fig. 2J). Overall, CRD tumors exhibited more aggressive organoids than LD tumors.

### Circadian desynchronization alters the TME

Breast TME is essential for the survival and immunosuppression of tumor cells (Binnewies, Roberts et al. 2018). We performed scRNA analysis on cells isolated from GEMM basal tumor tissues. Single cells were isolated from the tumors (Rodriguez de la Fuente, Law et al. 2021) without surface marker selection. Using proper QC and filtration, we obtained single-cell transcriptomic data for the studied tumors (N= 2/set). To analyze different cell populations in the TME, 5,804 QC-positive single cells (LD and CRD tumors) were clustered according to their expression profiles (Fig. 3A), and 20,764 genes were identified. The Seurat package was used for preliminary clustering. Cells were clustered based on specific cell markers. Using the established expression markers, such as for endothelial cells (*KLF2*)(Sangwung, Zhou et al. 2017), M2 macrophages (*SLC2A1*)(Wang, Wang et al. 2023), TNBC cancer cells (*TRPS1*)(Ai, Yao et al. 2021), proliferating cells (*TOP2A)*(Neubauer, Wirtz et al. 2016), myeloid cells (*FTH1*)(Moreira, Silva et al. 2022), progenitor cells (*NES*)(Bernal and Arranz 2018), adipocytes (*CLEC11A*)(Merrick, Sakers et al. 2019), mesenchymal cells (*S100A4*)(Fei, Qu et al. 2017), angiogenic cells (*RPL35A*)(Wu, Al-Eryani et al. 2021), breast cancer stem cells *(FXYD3*)(Li, Nishimura et al. 2023), dendritic cells or DCs (*CD68*) (Strobl, Scheinecker et al. 1998), cancer-associated fibroblast or CAFs (*COL1A2*)(Han, Liu et al. 2020), epithelial cells (*EPCAM*)(Trzpis, McLaughlin et al. 2007), cycling cells (*CDK4*)(Baker, Poulikakos et al. 2022), and B-cells (*CD83*) (Grosche, Knippertz et al. 2020), we found that CRD altered different cell populations in the TME (Fig. 3B). This showed that CRD induced a higher number of M2-macrophage, TNBC cancer cells, cycling cells, adipocytes, CAFs and proliferating cells but a lower number of B-cells, and DCs in the tumor (Fig. 3B). We next analyzed putative ligand–receptor (L-R) interactions in LD and CRD tumors and modeled cell–cell interactions between different cell types that play vital roles in maintaining cellular homeostasis. Cellular communication was investigated using CellChat, which shows the differences in the total number of L-R interactions between different cell types in the tumors (Fig. 3C). For example, the interaction between CAF with proliferating cells, TNBC cells, and M2 macrophages increased under CRD conditions compared to LD conditions (Fig. 3C). We identified 3429 and 3992 critical ligand–receptor pairings in LD and CRD tumors, respectively (Fig. S5A). Ligand–receptor interactions with increased and decreased strength were also identified (Fig. S5B). CellChat also identified signaling pathways linked to ligand–receptor interactions (Fig. 3D). Specific signaling pathways, including those involving fibroblast growth factor (FGF), WNT, and platelet-derived growth factor (PDGF), were exclusively detected in CRD tumors. Oncostatin M, Tumor necrosis factor (TNF), and NOTCH, were detected solely in LD tumors, and signaling pathways, including transforming growth factor β (TGFβ), epidermal growth factor (EGF) and fibronectin 1, were detected in both LD and CRD tumors (Fig. 3D). Gene Ontology (GO) analysis (Fig. 3E) indicated that genes associated with the acetyl CoA metabolic process, antigen processing (and presentation), nuclear division, regulation of cellular pH, carboxylic acid catabolic process, chromosome segregation were upregulated in CRD.

**Fig. 3.**
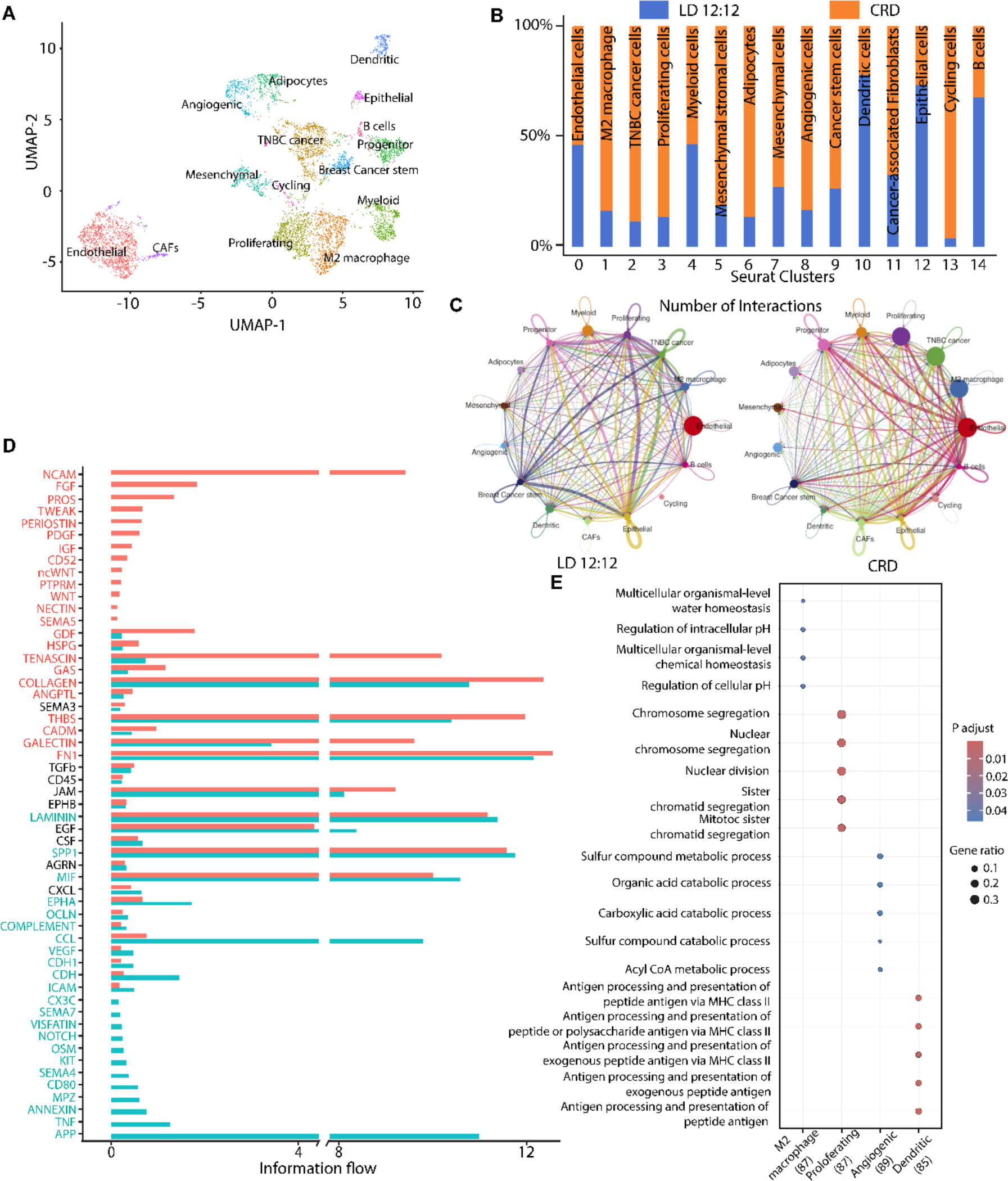
CRD alters the tumor microenvironment. Cell clusters from 10x Genomics scRNA-seq analysis visualized using Uniform Manifold Approximation (UMAP). Colors indicate clusters of various cell types **(A)**. Fraction of cells from tumors (LD and CRD) in each cluster **(B)**. Clusters were annotated for their cell types as predicted using canonical markers and signature-based annotation with Garnett. Expression markers for endothelial cells (*KLF2*), M2 macrophages (*SLC2A1*), TNBC cancer cells (*TRPS1*), proliferating cells (*TOP2A)*, myeloid cells (*FTH1)*, progenitor cells (*NES*), adipocytes (*CLEC11A*), mesenchymal cells (*S100A4*), angiogenic cells (*RPL35A*), breast cancer stem cells *(FXYD3*), dendritic cells (*CD68*), cancer-associated fibroblast (*COL1A2*), epithelial cells (*EPCAM*), cycling cells (*CDK4*), and B-cells (*CD83*). The comparison of the total number of interactions among different cell populations between LD and CRD tumors is shown in **C**. Edge width is proportional to the number of interactions, which assesses the number of ligand–receptor pairs contributing to the communication between two interacting cell populations. CellChat analysis quantitatively analyzes intercellular communication networks **(D)**. The analysis shows differential ligand–receptor interactions between LD and CRD tumors. Significant signaling pathways were ranked based on differences in the information flow within the inferred networks between LD and CRD tumors. Dot plot profiling of the Gene Ontology (GO) analyses in each cluster **(E)**. The dot plot represents the average expression of GOs per cluster. The color gradient of dots represents the expression level, whereas the size represents the percentage of cells expressing any genes per cluster. The number indicates the total number of genes identified belonging to GO pathways significantly enriched in the cluster.

### CRD induces immunosuppressive TME and turns tumors “cold”

The TME is critical for tumor progression and metastasis. Tumors are “hot” when they show signs of inflammation characterized by an infiltration of T cells mobilizing to fight the cancerous cells, whereas nonimmunogenic “cold” tumors lack infiltrating T cells, which makes it challenging to provoke an immune response with immunotherapy drugs (Ren, Guo et al. 2022). To investigate whether the increased prevalence of metastasis in CRD mice resulted from changes in the TME, we characterized different immune cell populations in the TME of LD and CRD tumors using scRNA-seq, flow cytometry, and immunostaining (MxIF). We determined whether CRD could influence the proportion of various immune cells in the TME and, in turn, promote tumor progression and metastasis. scRNA-seq with GEMM tumors showed that CRD reduced cytotoxic T-cell infiltration (Fig. S6A) and enhanced regulatory T-cell (T_reg_) infiltration (Fig. S6B). CRD-induced M2-macrophage populations while decreasing M1-macrophage populations (Fig. S6C). Flow cytometry analysis with CRD-induced 4T1 tumors supported the scRNA seq data. Our study showed a significant inhibition of the number of infiltrating CD8^+^ T cells (cytotoxic T-cells) (Fig. 4A) and an increase in the CD4/CD8 ratio (Fig. 4B) in CRD-induced tumors. The increased CD4/CD8 ratio in CRD-induced tumors also indicates an immunosuppressive microenvironment. FACS showed significant enrichment of the immunosuppressive T_reg_ (CD4^+^FoxP3^+^) population (Fig. 4C) in the CRD-induced tumors. We identified anti-tumor (M1) and pro-tumor (M2) macrophage populations in the LD and CRD-induced tumors. CRD induced M2-macrophage infiltration while reducing M1-macrophage infiltration (Fig. 4D). Tumors from mice under CRD conditions exhibited a ratio (M1:M2) that favored M2-mediated immunosuppression (Fig. 4E) and facilitated tumor growth. CRD significantly enhanced the CD45^+^ immune cell population in tumors (Fig. 4F). CRD-induced tumors showed a lower dendritic cell population compared to LD-induced tumors which support the anti-cancer properties of dendritic cells (CD11c) (Fig. 4G). Flow cytometry study also showed a significant increase in the MDSC population (CD11b) (Fig. 4H) in CRD-induced tumors, which further supports an immunosuppressive microenvironment. A donut chart shows the differences in different immune cell populations in LD vs CRD-induced tumors (Fig. 4I). Our immunofluorescence assay showed that CD8+ T-cell infiltration was decreased, but M2 macrophages (CD163) and FOXP3 (T_reg_) expression was enhanced in CRD-induced GEMM tumors compared with those in LD GEMM tumors (Fig. 4J), making the tumor “cold.” CRD increased EMT gene expressions, such as fibronectin 1 (FN1) and Snail (SNAIL 1) (Fig. 4K). There was an increase in S100A4^+^ and α-smooth muscle actin (α-SMA^+^) CAF populations in CRD-induced tumors compared to those in LD tumors (Fig. 4L).

**Fig. 4.**
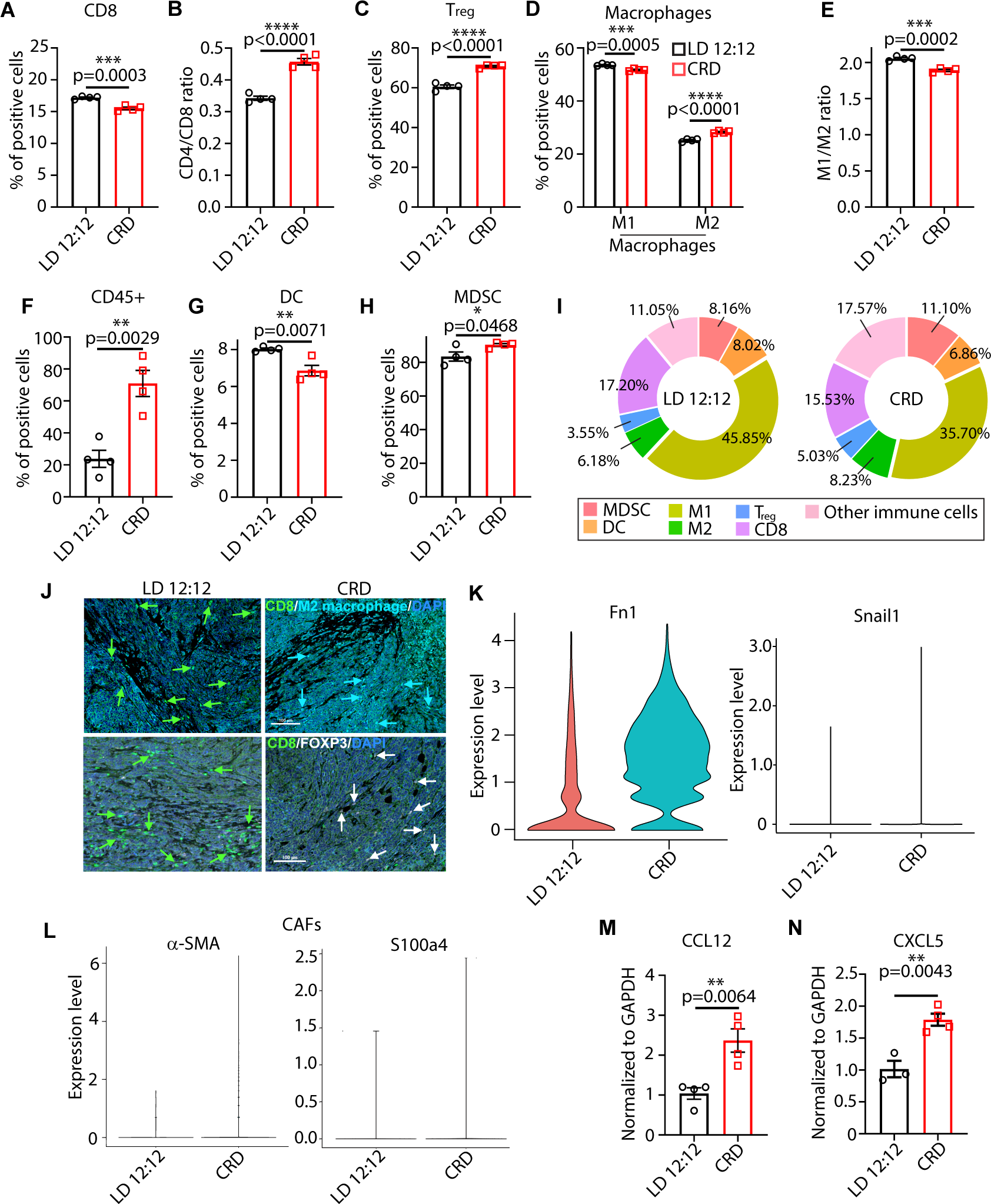
CRD turns tumors “cold.” Flow cytometry showed a significant decrease in cytotoxic CD8 T cell population (A). An overall ratio of CD4/CD8 is shown in **(B)**. Flow cytometry revealed a significant increase in the FoxP3^+^ T_reg_ population in the tumors of CRD mice (n = 4) **(C).** The percentage of antitumor M1 macrophage population and pro-tumor M2 macrophage population (n = 4) in LD and CRD-induced 4T1 tumors is shown in **(D),** and an overall ratio of M1/M2 is shown in **(E)**. The percentage of total leukocytes (CD45^+^ cells) in LD and CRD-induced tumors is shown in **(F)**. The percentage of DC cells **(G)** and MDSC **(H)** is shown in LD and CRD-induced tumors via flow cytometry (n=4). P-value obtained from an unpaired two-sided *t*-test. The relative distribution of key immune cell types in LD and CRD-induced tumors is represented in the donut chart **(I).** Representative immunostaining image of cytotoxic CD8 T-cell (CD8, green), M2 macrophages (CD186, Cyan), and T_reg_ (FOXP3, White) in the LD and CRD-induced GEMM tumors (n = 4) Scale bar= 100μm **(J)**. Single-cell RNA seq (scRNA Seq) study showed epithelial to mesenchymal (EMT) markers (Fibronectin and Snail1) **(K),** and cancer-associated fibroblast (CAF) population based on S100a4, and α-SMA markers **(L),** in LD and CRD-induced tumors. The expression of chemokines *Ccl12* **(M)** and *Cxcl5* **(N),** is shown using real-time PCR in LD and CRD-induced tumors (n = 3 - 4).

Our data suggest that CRD enhances tumor progression and lung metastasis by enhancing the immunosuppressive microenvironment and increasing the CSC population. Our scRNA-seq, and immunostaining results confirmed the “cold” characteristics of CRD tumors. The gating strategy used for flow cytometry is shown in Fig. S7. Chemokines play a pivotal role in the migration pattern of immune cells into the tumor and facilitate the immunosuppressive TME by enhancing the differentiation and infiltration of immunosuppressive cells, such as T_reg_ cells, myeloid-derived suppressor cells (MDSCs), and tumor-associated macrophages (TAMs). To investigate whether CRD altered the chemokine-cytokine network in the studied mice, we quantified the levels of 40 circulating chemokines in the plasma of LD and CRD mice using a Proteome Profiler. Marked increases in circulating IFN-γ, IL-1β, G-CSF, and IL-16 levels were observed in CRD mice (Fig. S8A). As plasma cytokine/chemokine levels do not represent the TME, we used the data from our scRNA-seq and real-time PCR to assess chemokine/cytokine network expression levels in LD and CRD tumors. Levels of specific cytokines/chemokines known to favor an immunosuppressive microenvironment (Vilgelm and Richmond 2019), such as CCL12 (Fig. 4M) and CXCL5 (Fig. 4N), were upregulated in CRD-induced GEMM tumors. Levels of CCL28, known to induce T_reg_ infiltration and create an immunosuppressive environment (Yang 2017), were upregulated in CRD tumors, as shown using scRNA-seq (Fig. S8B), but not significantly, as shown using real-time PCR (Fig. S8C). Levels of IL-17β and IL10, were upregulated modestly (but not significantly) in CRD-induced 4T1 tumors (Fig. S8C).

### Malignant human breast cancer cells have high CRD

Earlier studies (Hadadi 2020) and the present study used GEMMs to investigate the effect of CRD on tumorigenesis. However, more insights are needed into the molecular mechanisms underlying CRD at the single-cell level and in human tumors. To investigate the process underlying CRD during mammary tumorigenesis, we used a scRNA sequence dataset from a previously published study (Wu, Al-Eryani et al. 2021). The dataset included TNBC, HER2-positive, and luminal tumors. Using large-scale copy number variation (CNV) from the transcriptomics data (Tirosh, Izar et al. 2016), we separated definitive malignant cells from potential nonmalignant cell populations in TNBC (Fig. 5A), luminal (ER^+^/PR^+^) (Fig. S9A), and HER2 tumors (Fig. S9B). Based on a previous study (He, Fan et al. 2022), the CRD score was calculated using the expression of 358 circadian-related genes (He, Fan et al. 2022). Next, we quantified the CRD scores of malignant (and nonmalignant) cells in different types of human breast tumors. Aggressive TNBC comprised significantly (P < 0.01) more malignant cells with high CRD scores (Fig. 5B) than luminal tumors (Fig. S9C). Malignant cells in HER2 tumors also showed a high CRD score (Fig S9D). *Cry1*, *Per1*, and *Per2* expression decreased in TNBC malignant cells compared to non-malignant cells, as shown using the scRNA-seq dataset (Fig. 5C). To define the role of CRD in large patient populations, we used publicly available RNA-seq data from tumors (N=1104 patients) and surrounding tissues (N = 113) of patients with breast cancer in The Cancer Genome Atlas (TCGA). The expression of core clock genes (*Per1*, *Per2*, *Per3*, *Cry2*, *Rorc*, *Rorb*, *Nr1d1*, and *Arnt2*) was significantly (P < 0.05) lower in the tumors than in the surrounding tissues (Fig. 5D). These data suggest that the core clock is disrupted in human breast tumors.

**Fig. 5.**
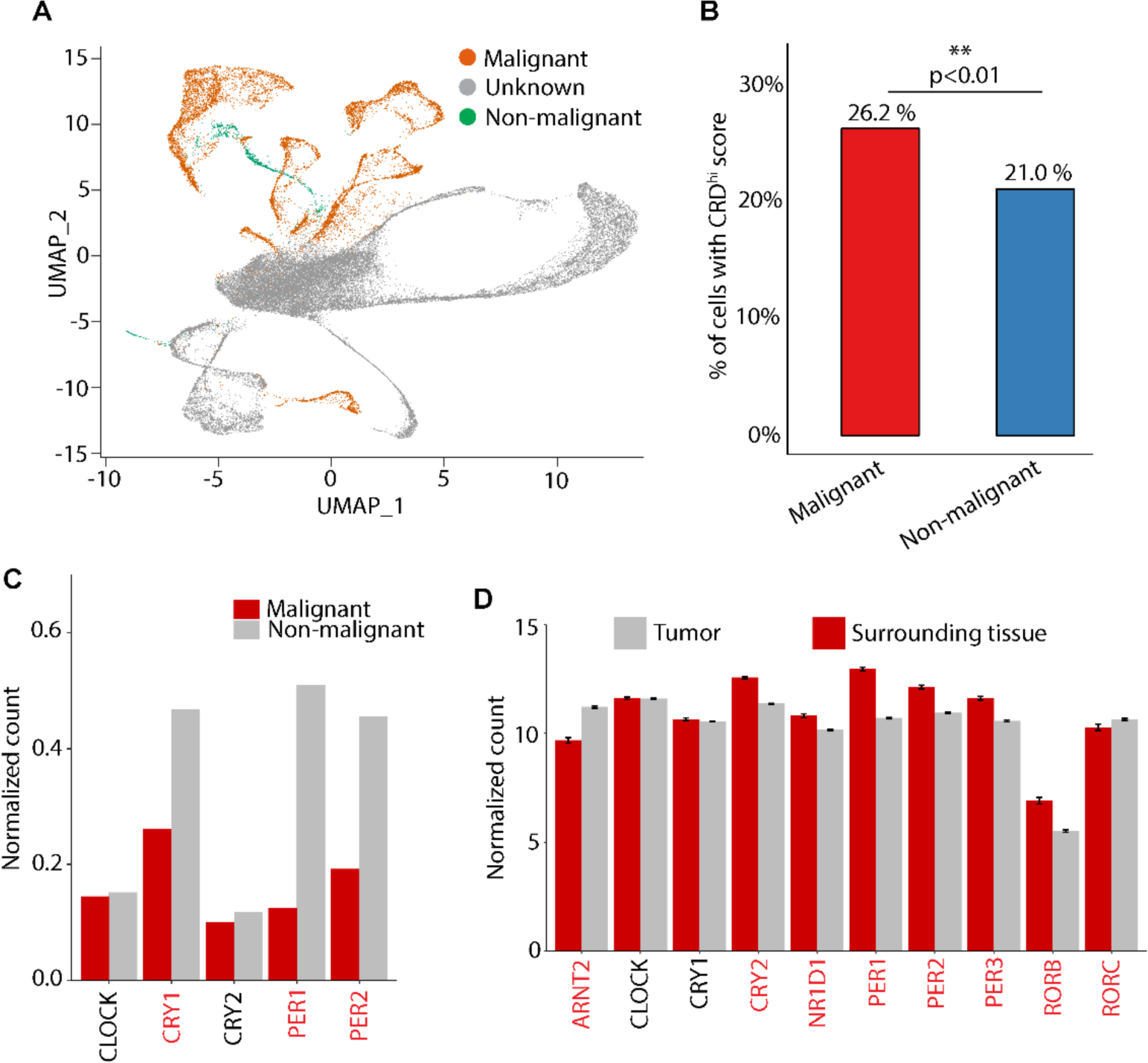
The human circadian clock is disrupted in breast cancer. UMAP of malignant and nonmalignant cells (total cells = 42,512) captured across patients with (n = 9) triple-negative breast tumors. Moving from the plot in **A**, 30,719 cells had an unknown inferCNV call and 11,793 had a known inferCNV call. These 11,793 single cells were used for downstream analysis to detect “malignant” and “nonmalignant” cells. The percentage of malignant and nonmalignant cells with CRD^high^ is shown in **B**. The mean normalized expression values of the core circadian genes by the cell type (malignant and nonmalignant) are presented in **C**. Red represents significant changes in expression between malignant and nonmalignant cells. **(D)** The mean expression of the clock genes (+/- standard error of mean) over patients by the type of tissue analyzed in TNBC tumors (n = 1104) and surrounding tissue (n = 113) using TCGA.

### CRD creates “cold” tumors by inducing LILRB4 expression

Next, to identify the molecular mechanism that regulates the CRD-induced immunosuppressive TME, we searched for transcripts differentially expressed between the samples (mammary tumors and glands) collected from mice exposed to ordinary light and dark conditions (LD 12:12) and mice subjected to CRD. Differential gene expression (DGE) analysis of our scRNA-seq data revealed that LILRB4 was upregulated in the tumor samples from CRD-induced GEMMs (Fig. 6A, S10A). LILRB4, an immunoreceptor tyrosine-based inhibitory motif-containing receptor, suppresses T-cell activity via a signaling pathway involving apolipoprotein E (APOE), SH2-domain-containing protein tyrosine phosphatase-2 (SHP-2), urokinase-like plasminogen activator (uPAR or Plaur), and arginase 1 (ARG-1) (Sharma 2021). Recent studies have shown that LILRB4 creates an immunosuppressive microenvironment in AML (Deng, Gui et al. 2018) and solid cancers (Sharma 2021) by inhibiting CD8^+^ T-cell infiltration and inducing T_reg_. Real-time PCR showed that CRD enhanced LILRB4 levels significantly in the CRD-induced GEMM tumors (Fig. 6B), and CRD-induced 4T1 tumors (Fig. 6C). The immunofluorescence analysis showed an elevation of ARG1 expression in CRD-induced GEMM tumors (Fig. 6D). The transcript level of ARG1 was significantly increased in CRD-induced tumors (Fig. 6E). Increased LILRB4 expression enhanced the immunosuppressive TME as CD8+ T cell infiltration decreased, whereas M2-macrophage (CD163) and T_reg_ (FOXP3) populations increased in CRD-induced mammary tumors from GEMMs, as shown using immunofluorescence assay (Fig. 6F). Furthermore, LILRB4 transcript levels were significantly increased in CRD-induced mammary glands (Fig. 6G). Mammary gland LILRB4 transcript levels exhibited robust diurnal oscillations, increasing during the light phase under normal and dark conditions. Under CRD, the amplitude of this rhythm was dampened, but the phase was reserved (or only slightly shifted), and the transcript level peaked during the dark phase (Fig. S10B). The transcript level of ARG1 increased modestly but non-significantly in the CRD-induced mammary glands (Fig. S10C). The transcript level of the chemokine CCL12 increased significantly in CRD-induced mammary glands (Fig. S10D). The transcript level of CXCL5 is raised modestly but not significantly in CRD-induced mammary glands (Fig. S10E). Overall, these changes in the immune milieus of the CRD-induced mammary microenvironment can decrease adaptive immunity and enhance susceptibility to cancer and other diseases.

**Fig. 6.**
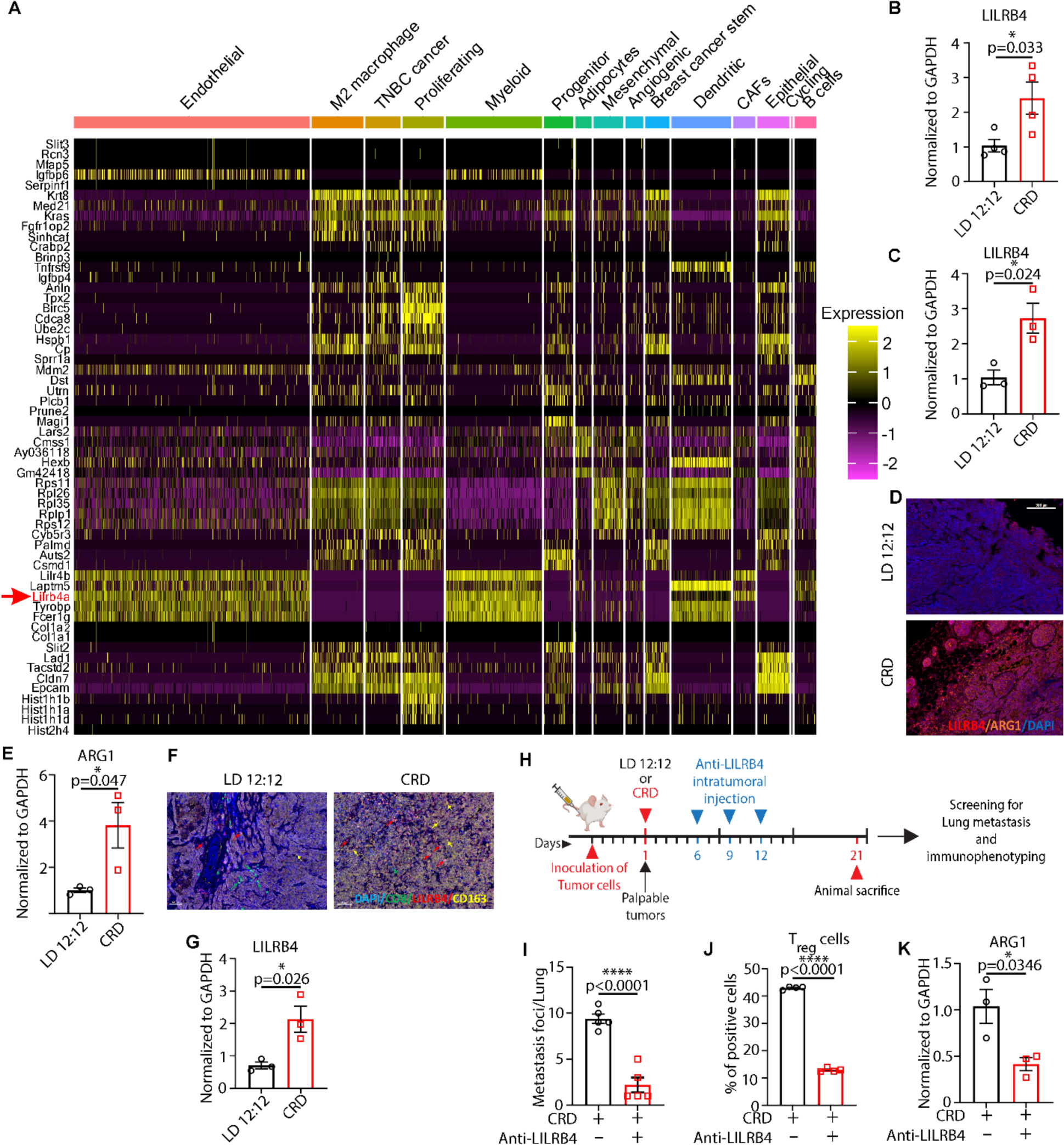
CRD enhances LILRB4a expression to create an immunosuppressive tumor microenvironment. **(A)** Heatmaps of differentially expressed genes between LD 12:12 and CRD tumors identified using DESeq2 analyses of scRNA-seq. The transcript level of LILRB4 was analyzed in LD and CRD-induced GEMM tumors **(B)** and 4T1 tumors **(C)** using real-time PCR. **(D)** The expression of arginase 1 (*Arg1*, a downstream target of LILRB4a) was analyzed using immunofluorescence. Scale bar: 300μm **(E)** The transcript level of *Arg1* in LD and CRD-induced GEMM tumors was analyzed using real-time PCR. **(F)** Representative MxIF showing the expression of LILRB4, CD8, CD163, and FOXP3 markers in LD and CRD-induced GEMM tumors. Scale bar: 50μm. **(G)** The transcript level of *Lilrb4* in LD and CRD-induced mammary glands was analyzed using real-time PCR. **(H)** 4T1 cells were injected into the mammary fat pads of BALB/cJ mice for the LILRB4-targeted immunotherapy. The number of metastatic foci in the lungs of CRD (control) and CRD (LILRB4-antibody) mice at the time of their sacrifice is shown in **I.** The percentage of T_reg_ cells in control and LILRB4-antibody-treated CRD-induced tumors is shown in **J.** The transcript level of *Arg1* in control and LILRB4-antibody-treated CRD-induced tumors is shown by real-time PCR **(K).**

To investigate whether LILRB4 expression was responsible for creating an immunosuppressive TME, we explored the effects of LILRB4-immunotherapy on tumor progression in CRD mice by treating them with an LILRB4 antibody (Sharma 2021). Briefly, 4T1 cells were orthotopically transplanted into BALB/cJ mice to develop TNBC tumors. Four to five days (∼4-5) days after the transplantation, once palpable tumors developed, mice were placed into LD or CRD condition. The LILRB4 antibody was administered on days 6, 9, and 12 after developing palpable tumors. Mice were sacrificed 3 weeks after the development of palpable tumors (day 21) (Fig. 6H), and lung metastasis and the TME were examined. We observed a significant decrease (P<0.0001) in the prevalence of lung metastasis in LILRB4-antibody treated mice under CRD condition (Fig. 6I). Flow cytometric analysis showed an inhibition of the T_reg_ cell population in LILRB4-antibody treated tumors under CRD condition, which alleviate the immunosuppressive microenvironment (Fig. 6J). LILRB4-targeted therapy inhibited ARG1 transcript level (Fig. 6K) in CRD-induced tumors. We also observed a modest but non-significant decrease in CCL12 (Fig. S10F) and CXCL5 (Fig. S10G) transcript levels in CRD-induced tumors. No significant difference was observed in the number of lung metastasis foci in LD mice treated with the LILRB4-antibody (Fig S10H). LILRB4-antibody treatment under LD conditions does not decrease the T_reg_ population (Fig. S10I), ARG1 transcript level (Fig. S10J), and the chemokines (CCL12, CXCL5) (Fig. S10K, L) significantly. These findings fortify the notion that elevated LILRB4 contributes to CRD-induced aggressive tumorigenesis.

## Discussion

Following the chronic jet lag protocol, here, we showed that CRD interrupted the biological clock, affected the mammary gland morphology, and accelerated aggressive basal mammary tumorigenesis and lung metastasis by altering the TME. CRD induced an immunosuppressive TME and turned the tumors “cold.” Malignant human breast cancer cells were shown to have high CRD scores. We also identified the molecular mechanisms underlying CRD-induced aggressive mammary tumorigenesis: CRD created “cold” tumors by inducing LILRB4 expression.

CRD-induced mammary glands exhibited underdeveloped (and abnormal) TEBs and reduced branching morphology. CRD-induced mice showed mammary gland ductal hyperplasia, which can be the early sign of ductal cancer (Behbod, Gomes et al. 2018), as seen by H&E and Ki67 staining. This indicates the role of the circadian clock in mammary gland development and the potential role of CRD in breast cancer initiation. Our study showed that CRD accelerated spontaneous mammary tumor initiation and lung metastasis but not tumor progression, indicating that CRD affects the early events of tumor initiation, rather than the growth of the established tumors. LILRB4 expression was increased in mammary glands and CRD-induced mammary tumors, revealing an unidentified mechanism responsible for CRD-induced tumorigenesis, lung metastasis, and mammary gland development. The jet lag protocol used here mimics the effects of rotating shift work or frequent eastbound trans-meridian flights and has been previously shown to cause severe perturbations of circadian rhythmicity (Papagiannakopoulos, Bauer et al. 2016, Hadadi, Taylor et al. 2020, Pariollaud, Ibrahim et al. 2022). Consistent with the results of published studies (Pariollaud, Ibrahim et al. 2022), we found that CRD affected the rhythmicity of many clock genes (e.g., *Clock, Bmal1, Per1*, and *Cry1*) in the mammary glands; however, *Per2* expression remained rhythmic after CRD, indicating that *Per2* is more sensitive to phase entrainment signals of molecular oscillators in peripheral tissues.

Using scRNA-seq and TCGA data, we showed that the expression of certain circadian genes was decreased in aggressive human TNBCs compared with that in the surrounding mammary tissues. Our analysis also revealed that the malignant cells in aggressive human TNBCs had higher CRD scores and downregulated *PER1*, *PER2*, and *CRY1* significantly than nonmalignant cells.

Our study showed that CRD induced an immunosuppressive tumor microenvironment, which could be involved in increased tumor burden in mice maintained under the CRD schedule. The TME consists of natural killer cells, CD8^+^ T cells, CD4^+^ helper T (T_h_) cells, proinflammatory macrophages (M1), and DCs, which elicit antitumor immune responses, whereas the presence of MDSCs and FOXP3^+^ T_reg_ counteracts tumor immunity (Labani-Motlagh, Ashja-Mahdavi et al. 2020). Our scRNA-seq study, and FACS showed that CRD decreased cytotoxic T-cell infiltration, and the DC population. This supports the recent finding that DC and T-cell-autonomous circadian clocks are responsible for time-of-the-day-dependent antitumor effects (Wang, Barnoud et al. 2023). CRD-induced S100a4^+^ CAF, and α-SMA^+^ CAF populations are known to promote tumor progression and metastasis and contribute to an immunosuppressive TME (Costa, Kieffer et al. 2018). We found an increase in the M2 macrophage, and MDSC population and a decrease in the M1 macrophage population in CRD tumors. M2 macrophages play a significant role in creating an immunosuppressive microenvironment (Hao, Lu et al. 2012), whereas M1 macrophages induce inflammation and kill tumor cells (Hao, Lu et al. 2012, Pan, Yu et al. 2020). MDSCs are also known to create an immunosuppressive environment by suppressing T-cell responses and CD8^+^ T-cell infiltration (Li, Shi et al. 2021). Our FACS results corroborated the scRNA-seq results and confirmed that CRD enhanced the T_reg_ population. CRD increased the leukocyte (CD45^+^) population, corroborating that the jet lag schedule enhances leukocyte levels in melanoma (Aiello, Fedele et al. 2020). Our scRNA-seq and real-time PCR studies showed that CRD induced a pro-tumorigenic and immunosuppressive microenvironment by increasing the expression of CCL12 and CXCL5. The upregulation of CCL12 (human orthologs of CCL2) (Yang J 2020) and CXCL5 reportedly lead to the accumulation of MDSCs (Li, Shi et al. 2021). CRD also enhances CCL28 production, which leads to the recruitment of T_reg_ cells in the TME (Ren, Yu et al. 2016).

We further demonstrated that CRD enhanced the expression of LILRB4 *in vivo* in healthy mammary glands and mammary tumors. Recent studies have shown that LILRB4 creates an immunosuppressive microenvironment in AML (Deng, Gui et al. 2018) and solid cancers (Sharma 2021) by inhibiting CD8^+^ T-cell infiltration and inducing T_reg_. LILRB4 inhibited the differentiation of CD8^+^ and CD4^+^ T cells (Deng, Gui et al. 2018). By activating the JAK/STAT signaling pathway, LILRB4 controls the expression of cytokines in macrophages (Truong, Hong et al. 2019). In different malignant cancers, LILRB4 expression in MDSCs is associated with decreased survival in patients (de Goeje, Bezemer et al. 2015). A recent study showed that LILRB4 blockade increased the proportions of tumor immune infiltrates, effector (T_eff_) levels, and altered the TME toward reduced immunosuppression in solid tumors (Sharma 2021). Together, these findings reveal that LILRB4 creates an immunosuppressive microenvironment, thereby decreasing its antitumor efficacy. Our study showed that CRD activated LILRB4 signaling in healthy mammary glands and aggressive TNBCs. Using real-time PCR and immunofluorescence analysis, we observed increased *ARG1* expression in CRD-induced tumors. Analysis of the TME using multiplexing immunostaining showed an immunosuppressive microenvironment as the cytotoxic T-cell and M1 macrophage populations decreased. In contrast, T_reg_, and M2 macrophage populations were increased in CRD tumors. We found that the disruption of circadian rhythm upregulated LILRB4 expression, which correlated with mammary tumor progression and abnormal morphology in the mammary glands.

CRD upregulated the expression of immunosuppressive chemokines (CCL12) in mouse mammary glands. CCL2 (an orthologous of mouse CCL12) increases stromal density and elevates cancer risk (Sun, Glynn et al. 2017). An immunosuppressive environment can increase susceptibility to opportunistic infections (Shu Kurizky, Dos Santos Neto et al. 2020) in CRD-induced patients. This study demonstrated how an inhibitory immune receptor altered the TME and influenced aggressive mammary tumorigenesis in response to CRD. We showed that the targeted LILRB4 immunotherapy reduced CRD-induced lung metastasis by inhibiting immunosuppressive TME.

Our findings suggest that LILRB4 is a potential therapeutic target for mitigating CRD-induced cancer risk in populations exposed to chronic CRD, such as shift workers. LILRB4 inhibition was also found to improve the immunosuppressive microenvironment. However, how the disruption of the cell-autonomous molecular clock by CRD could enhance LILRB4 expression remains unknown. Further investigations are needed to determine why and how CRD enhances LILRB4 expression. Additional investigations are required to determine whether the chronic elevation of LILRB4 levels in the mammary glands in response to CRD occurs early in the disease process and whether LILRB4 is present in other anatomical locations.

In conclusion, our study provides the link between CRD and aggressive basal mammary tumorigenesis, metastasis, and abnormal mammary gland morphology. Based on our data, we propose a signaling cascade (**Fig. 7)** as a possible mechanism behind CRD-induced mammary gland disruption, enhanced aggressive tumorigenesis, lung metastasis, and immunosuppressive TME.

**Fig 7.**
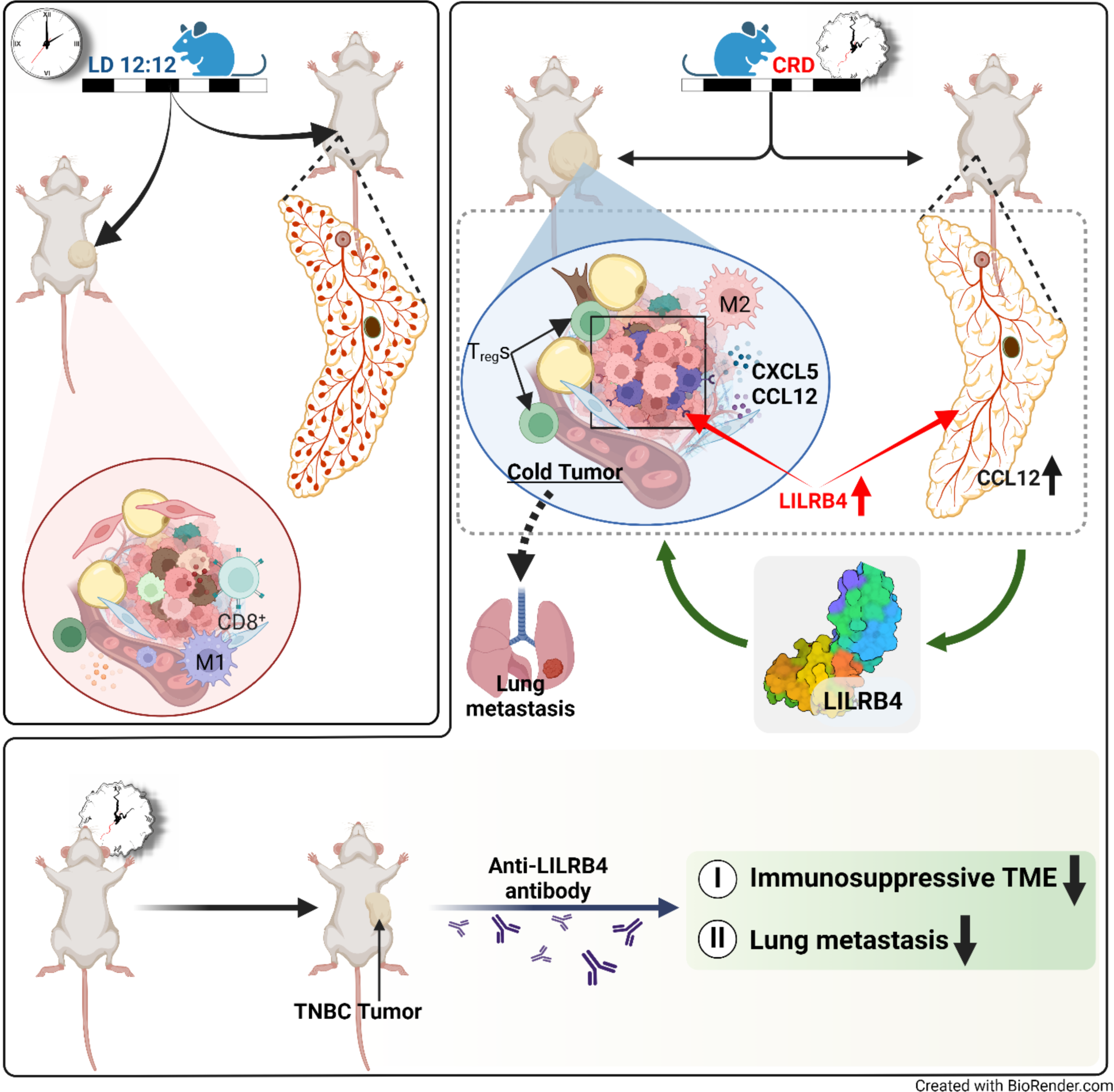
Diagrammatic representation of the molecular mechanism of CRD-induced immunosuppressive microenvironment in breast cancer. CRD enhances aggressive TNBC and lung metastasis by creating an immunosuppressive tumor microenvironment. CRD also disrupts the mammary gland morphology. Our study revealed that CRD enhanced LILRB4 expression, which induces an immunosuppressive “cold” TME by increasing the M2 macrophage and T_reg_ populations and decreasing M1 macrophage infiltration. Inhibition of LILRB4 via LILRB4-targeted antibody alleviates CRD-induced immunosuppressive TME and inhibits lung metastasis.

## Supporting information

OO_CRD-Suppl_MS_24

## List of abbreviations

LD: Light and Dark
CRD: Circadian Rhythm Disruption
LILRB4: leukocyte immunoglobulin-like receptor 4
TNBC: Triple-negative Breast Cancer
ER: Estrogen receptor
PR: Progesterone receptor
HER2: Human Epidermal Growth Factor Receptor 2
SCN: Suprachiasmatic nucleus
CLOCK: Circadian locomotor output cycles kaput
BMAL-1: Brain and muscle Arnt-like protein-1
PAS: Per-Arnt-Sim domain
PER: Period
CRY: Cryptochrome
TME: Tumor microenvironment
GEMM: Genetically engineered mouse model
FVB: Friend LeukemiaVirus B
TEB: Terminal end bud
H&E: Hematoxylin and Eosin
AGR: Albumin-to-globulin ratio
scRNA-seq: Single-cell RNA sequence
CD: Cluster of differentiation
KLF2: Krüppel-like factors2
SLC2A1: Solute Carrier Family 2 Member 1
TRPS1: Transcriptional Repressor GATA Binding 1
TOP2A: Topoisomerase II Alpha
FTH1: Ferritin heavy chain 1
NES: Nestin
CLEC11A: C-Type Lectin Domain Containing 11A
S100A4: S100 Calcium Binding Protein A4
RPL35A: Ribosomal Protein L35a
FXYD3: FXYD Domain Containing Ion Transport Regulator 3
COL1A2: Collagen Type I Alpha 2 Chain
EPCAM: Epithelial cellular adhesion molecule
CDK: Cyclin-dependent kinase
M1: Type1 macrophage
M2: Type2 macrophage
CAFs: Cancer-associated fibroblasts
L-R: Ligand-receptor
FGF: Fibroblast growth factor
PDGF: Platelet-derived growth factor
TNF: Tumor necrosis factor
NOTCH: Neurogenic locus notch homolog protein
TGFβ: transforming growth factor-beta
CoA: Coenzyme A
MxIF: Multiplex immunofluorescence
Treg: Regulatory T-cell
α-SMA: Alpha-smooth muscle actin
FN1: fibronectin 1
FACS: Fluorescence-activated cell sorting
FoxP3: forkhead box P3
CSC: Cancer stem cell
MDSCs: Myeloid-derived suppressor cells
TAMs: Tumor-associated macrophages IFN Interferon
IL: Interleukin
G-CSF: Granulocyte colony-stimulating factor
CCL2/12: Chemokine (C-C motif) ligand 2/12
CXCL5: C-X-C motif chemokine 5
PCR: Polymerase chain reaction
TCGA: The Cancer Genome Atlas
DGE: Differential gene expression
APOE: apolipoprotein E
SHP-2: SH2 domain-containing protein tyrosine phosphatase-2
uPAR: Urokinase-like plasminogen activator
ARG1: Arginase 1
AML: Acute Myeloid Leukemia
mIHC: multiplex immunofluorescence histochemistry
ARNT2: Aryl hydrocarbon receptor nuclear translocator 2
NR1D1: Nuclear Receptor Subfamily 1 Group D Member 1
RORB & C: RAR Related Orphan Receptor B and C
ITIMs: Immunoreceptor tyrosine-based inhibitory motifs
ITAMs: Immunoreceptor tyrosine-based activating motifs
APCs: Antigen-presenting cells
JAK/STAT: Janus kinase/signal transducers and activators of transcription

## Availability of data and materials

All data needed to evaluate the conclusions in the paper are present in the paper and/or the Supplementary Materials.

## Author contributions

Conceptualization: T.R.S. Methodology: O.O., M.S., A.C, D.J.B., T.N., D.F., Y.J-H., J.J and T.R.S. Investigation: O.O., M.S., A.C, D.J.B., T.N.,C.,N., T.R.S. Visualization: O.O, M.S., D.F., YJ-H., T.R.S. Supervision: T.R.S. Analysis of data: T.R.S., B.M., J.J., A.K. Writing—original draft: O.O., M.S., T.R.S. Review and editing: A.K., B.M., L.F and J.J.

## Acknowledgments

We thank Dr. Mahul Chakraborty for helping with real-time PCR, Dr. Gus Wright for helping with flow cytometry experiments and analysis, histology core facility (Texas A&M University) for histology work, Ashley Nguyen, David Arreola, Subiksha Sankar, and Aaryan Amin for helping with circadian cabinets.

