## Supplementary material for "Disruption of Circadian Clock Induces Abnormal Mammary Morphology and Aggressive Basal Tumorigenesis by Enhancing LILRB4 Signaling": OO_CRD-Suppl_MS_24

**Supplementary Figures:**

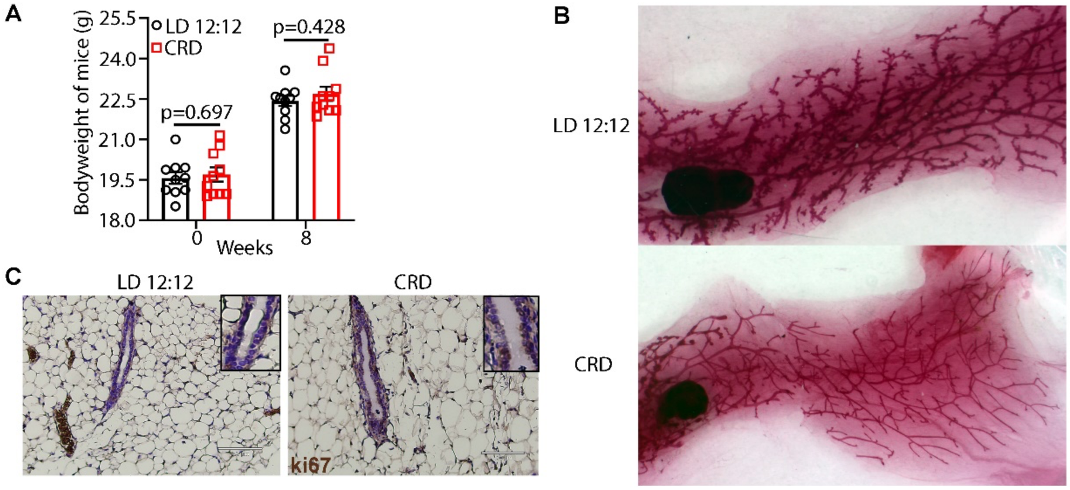


**Figure S1. The effect of CRD on mammary glands**. To investigate the effect of CRD on body weights, the body weights of the LD and CRD-induced mice were measured at week 0 and week 8 **(A).** The whole mount images of LD and CRD-induced mammary glands are shown in **B.** Staining for Ki67 revealed significant Ki67-positive cells in LD and CRD-induced mammary glands were stained for Ki67 marker **(C)**, scale bar: 100 μm.

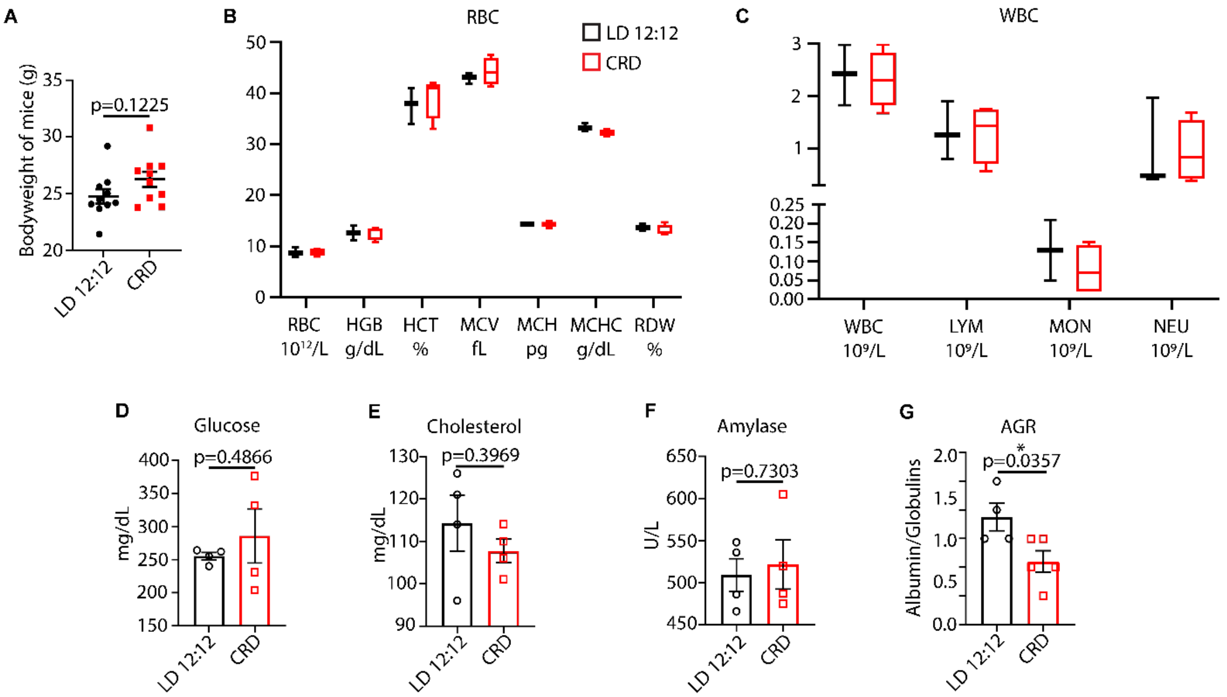


**Figure S2. CRD does not alter blood cell counts.** Female (FVB-Tg(C3-1-TAg)cJeg (C3-TAg) mice were housed in LD 12:12 or CRD condition and the weights were recorded at the end of the experiment **(A).** The total numbers of red blood cells (RBC) **(B)** and white blood cells (WBC) **(C)** in LD (n =5) and CRD (n =5) mice were counted. HGB, hemoglobin; HCT, hematocrit; MCV, mean corpuscular volume; MCH, mean corpuscular hemoglobin; MCH, mean corpuscular hemoglobin concentration; RDW, red cell distribution width; LYM, lymphocytes; MON, monocytes; NEU, neutrophils. Data presented as box-and-whisker plots. Variability is shown using medians (line in the box), 25th and 75th percentiles (box), and min to max (whiskers). The blood profile of some metabolites such as glucose (n=5 LD and n=5 CRD) **(D),** total cholesterol (n=5 LD and n=5 CRD) **(E),** amylase (n=5 LD and n=5 CRD) **(F),** and albumin to globulin ratio (AGR) (n = 5 LD and n = 5, CRD) **(G)** in LD and CRD-induced mice.

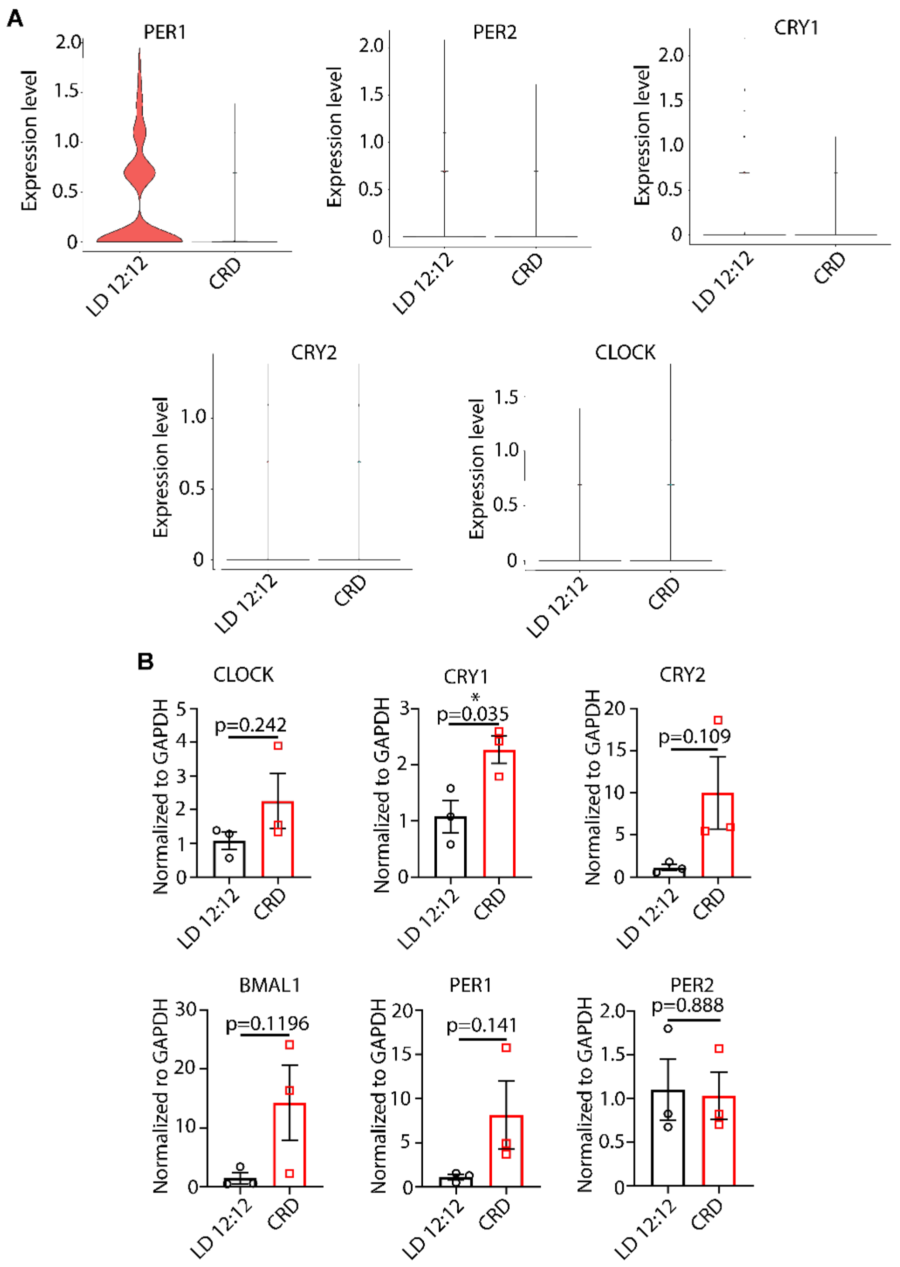


**Fig. S3. Core clock gene expression in tumors.** Expression of clock genes (PER1, PER2, CRY1, CRY2, CLOCK) in response to circadian rhythm disruption (CRD) is shown using scRNA-seq **(A)** and real time PCR **(B).**

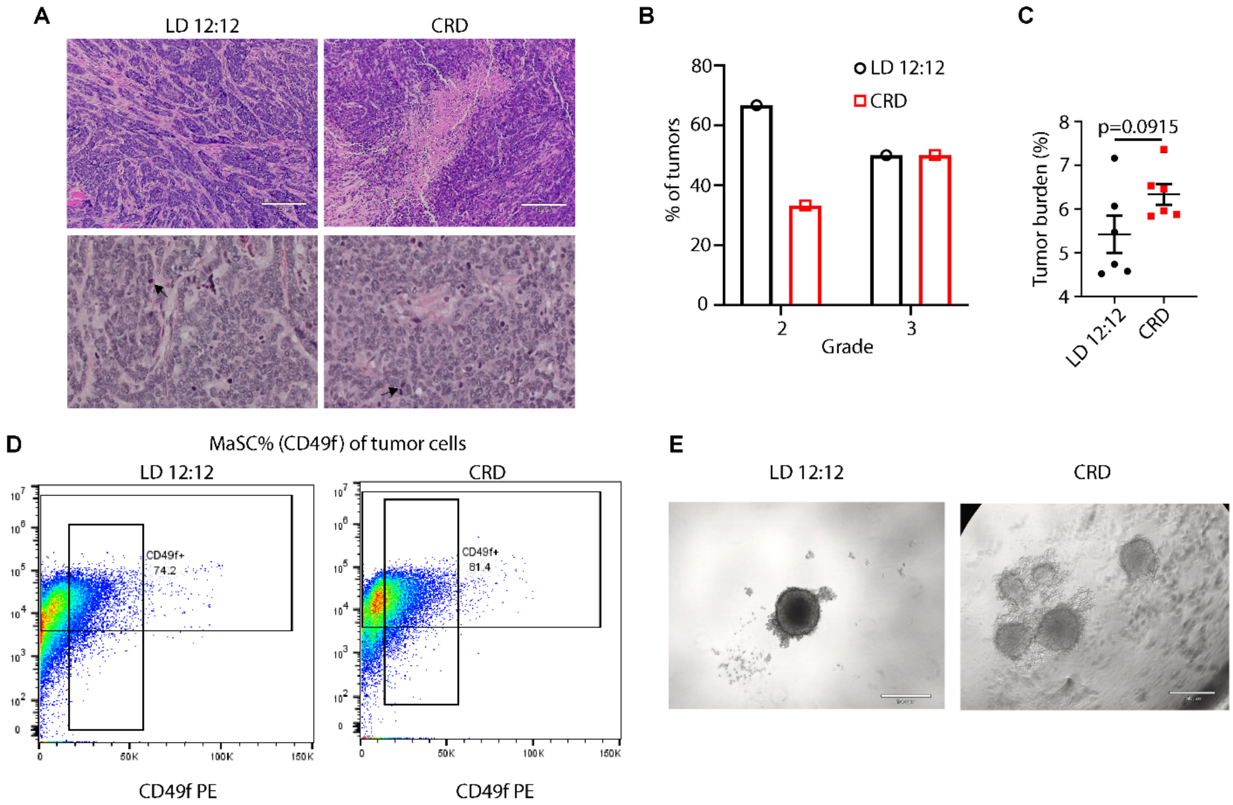


**Fig. S4**. **Histological analysis of LD and CRD-induced tumors.** H&E-stained tumors are shown **(A),** scale bar=150 μm and were analyzed and graded according to previous reports (Elston and Ellis, 2002). The data are presented as bar graph **(B).** The graphical representation of tumor burden in LD and CRD-induced 4T1 tumors **(C).** **(D)** CD49f marker profile prepared using flow cytometry. The mammosphere forming efficiency of LD and CRD tumor cells are shown in **E.** Scale bar: 150 μm.

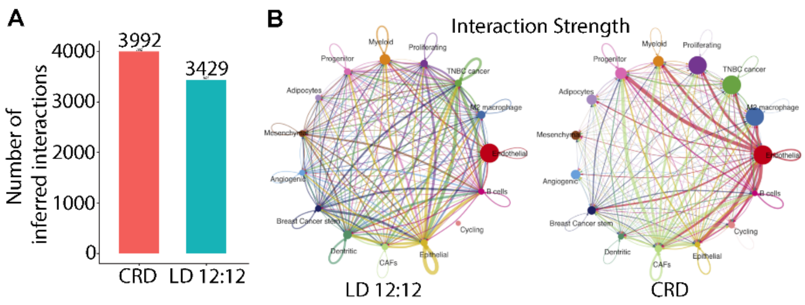


**Fig. S5. CRD alters the cell-cell interactions in tumor microenvironment.** CellChat analysis revealed 3429 and 3992 significant ligand-receptor interactions between cells in LD and CRD tumors, respectively **(A).** The interaction strength has been shown in **B.**

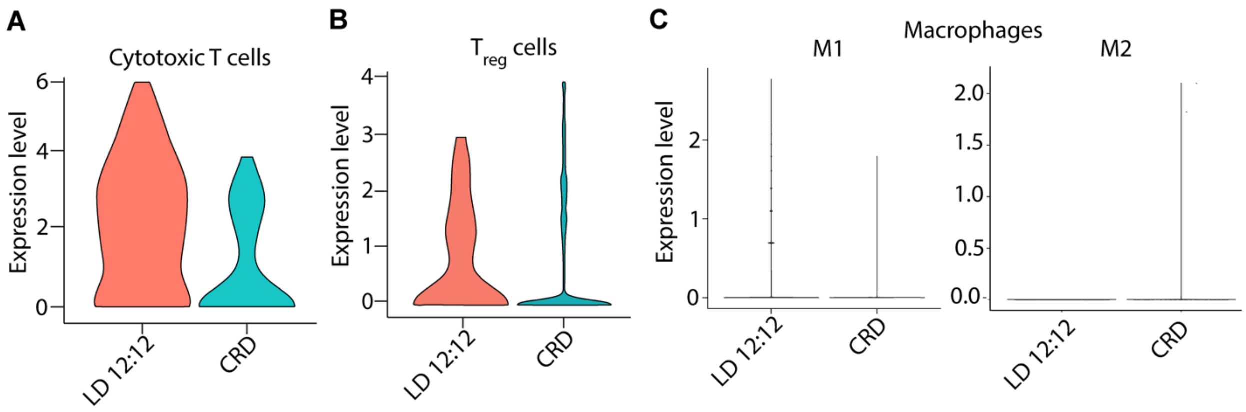


**Fig. S6. CRD alters the tumor immune microenvironment.** scRNA seq study showed that CRD reduced cytotoxic T cell population **(A)** and increased regulatory T cell population **(B).** CRD decreased M1 macrophage population and enhanced M2 macrophage population **(C).**

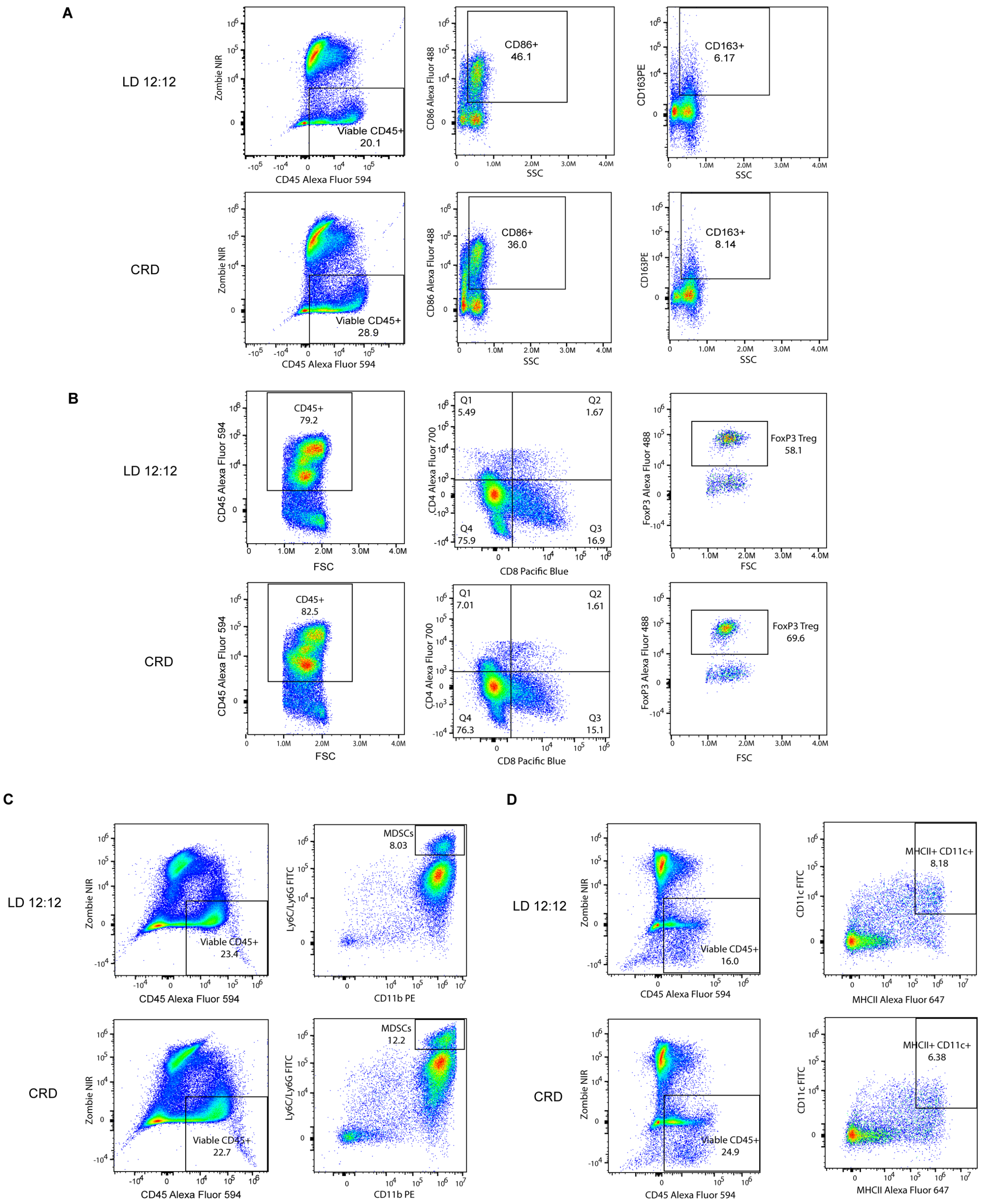


**Fig. S7. Flow cytometry gating strategy and immune cell numbers.** The gating strategy is used to identify CD45, macrophages (M1 and M2), DC, MDSCs, CD8T, and $T_{reg}$ (FOXP3) cells.

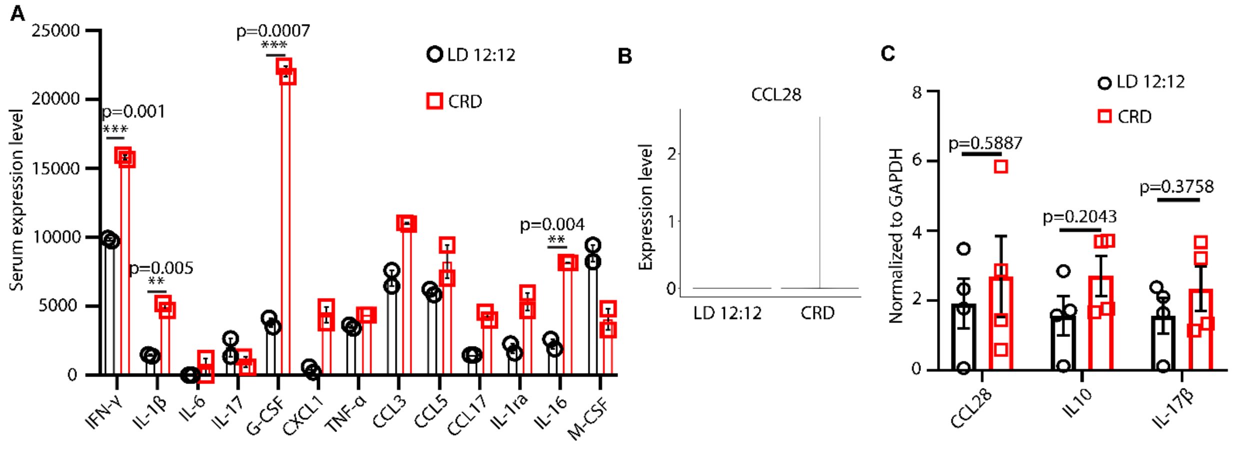


**Fig. S8. Cytokine and chemokine profile in LD and CRD mice. (A)** Proteome Profiler Mouse XL Cytokine Array kit was used to determine the circulating cytokine/chemokine concentrations in the CRD and LD mice. n=2 independent preparations for each condition **(A).** Quantitative estimation was performed by normalizing the densitometry value for each sample to that of a control. **(B)** CRD enhances CCL28 expression in mice, as shown by scRNA seq. **(C)** The transcript levels of CCL28, IL10, and IL-17β in LD and CRD-induced tumors were analyzed using real-time PCR.

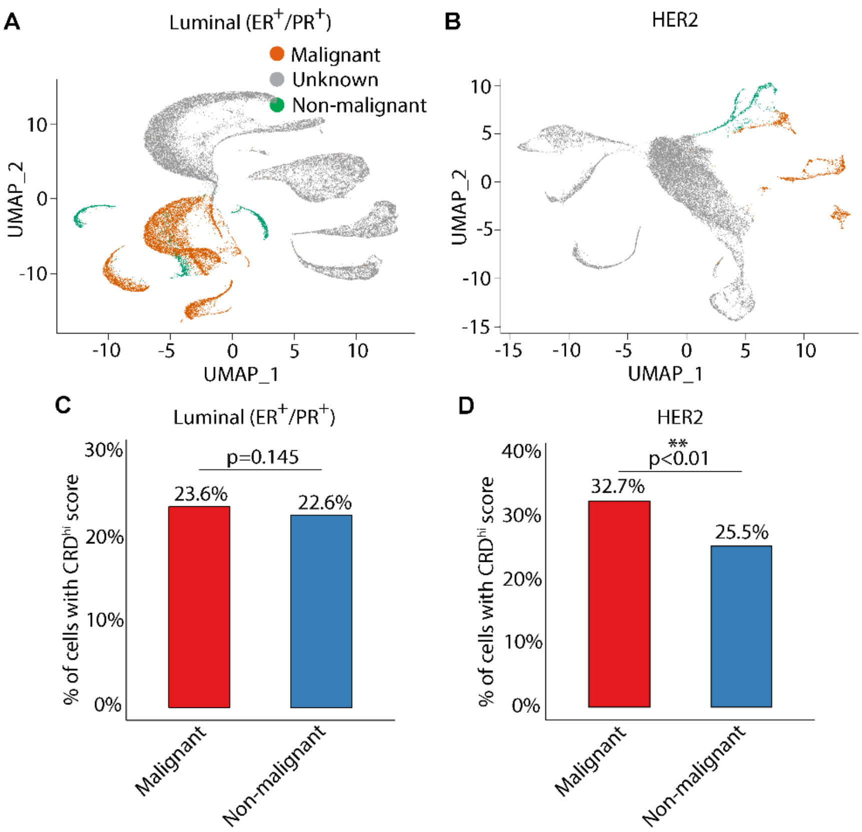


**Fig. S9. Circadian rhythm is disrupted in the malignant cells of the aggressive tumors**. UMAP of malignant and nonmalignant cells (total cells = 19,311) captured across the patients with luminal ($ER^+$ /$PR^+$) tumor **(A)** and UMAP of malignant and nonmalignant cells (total cells = 38,241) captured across the patients with $HER2^+$ tumor **(B).** The percentage of malignant and non-malignant cells with $CRD^{high}$ in $ER^+$ /$PR^+$ tumor **(C)** and $HER2^+$ tumor **(D)** is shown.

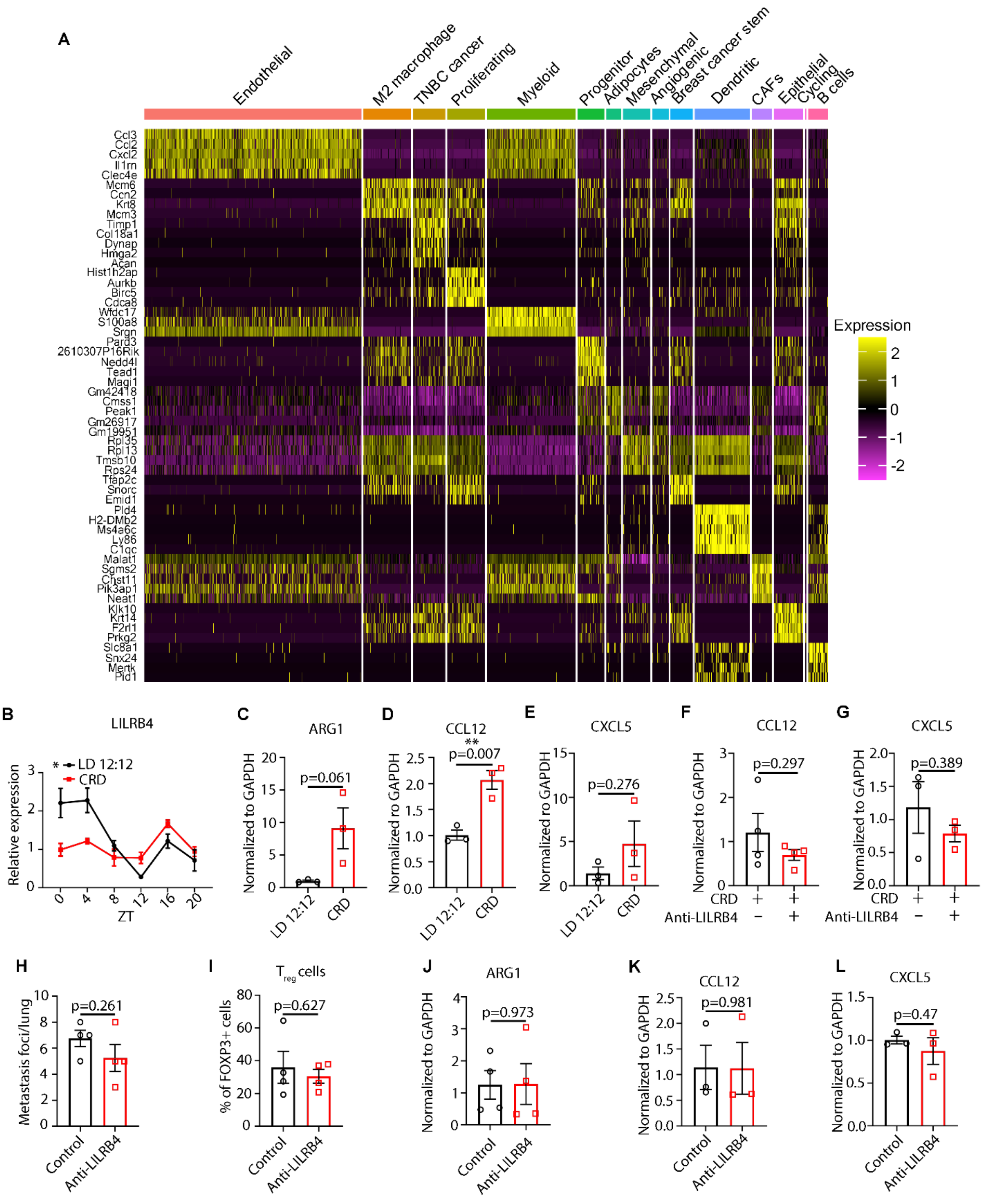


**Fig. S10. CRD induces mammary tumorigenesis by enhancing LILRB4 signaling pathway** **(A)** Heat maps of differentially expressed genes in LD tumors using DESeq2 analyses of scRNA-seq data. **(B)** Rhythmicity of the LILRB4 in the mammary gland was determined using JTK_Cycle analyses; **P*JTKcycle < 0.05, ***P*JTKcycle < 0.01, ****P*JTKcycle < 0.001, and *****P*JTKcycle

< 0.0001. **(C)** ARG1 transcript (target of LILRB4) level was determined in LD and CRD-induced mammary glands using real-time PCR. The transcript levels of CCL12 **(D)** and CXCL5 **(E)** were determined using real-time PCR**.** The transcript levels of CCL12 **(F),** and CXCL5 **(G)** were determined using real-time PCR in the LILRB4-targeted immunotherapy in CRD-induced tumors. The number of metastatic foci in the lungs of LD (control) and LD (LILRB4-antibody) mice at the time of their sacrifice is shown in **H.** The percentage of $T_{reg}$ cells in control and LILRB4-antibody-treated LD-induced tumors is shown in **I.** The transcript level of ARG1 in control and LILRB4-antibody-treated LD-induced tumors is shown by real-time PCR **(K).** The transcript levels of CCL12 **(K),** and CXCL5 **(L)** were determined using real-time PCR in the LILRB4-targeted immunotherapy in LD-induced tumors.

**Supplementary Data**

**Data S1: List of Primers**

| Genes | Strand | Sequence |
|---|---|---|
| GAPDH | Forward | 5'- AAC AGC AAC TCC CAC TCT TC -3' |
| | Reverse | 5'- CCT GTT GCT GTA GCC GTA TT -3' |
| CLOCK | Forward | 5'- CCA CCA CAG CAG TTC TTA CA -3' |
| | Reverse | 5'- TGC TCT GTT GTA GTG GAA AGG -3' |
| CRY1 | Forward | 5'- CTG AGG CAA GCA GAC TGA ATA -3' |
| | Reverse | 5'- CCC TCC ATT CCC ATT AGA GTT AG -3' |
| CRY2 | Forward | 5'- AGA GAC TAG TCG GCT CAA CA -3' |
| | Reverse | 5'- GGA AGG GAC AGA TGC CAA TAG -3' |
| PER1 | Forward | 5'- CAG TCA GAG CAG CCA TAC AA -3' |
| | Reverse | 5'- GTC CTG GAG CAC ACA CTT AAT -3' |
| PER2 | Forward | 5'- AAG CTG TCA CCA CCA TAG AAA G -3' |
| | Reverse | 5'- GCA AGG AGG CTG GTT CTT ATA G -3' |
| BMAL1 | Forward | 5'- CAT CAA GAC GAC ATA GGA CAC C -3' |
| | Reverse | 5'- CAT CGA CTT CGT AGC GTG ATA A -3' |
| gp49B (LILRB4a) | Forward | 5'- ATT CTG GAA CCC AGG ACT GC -3' |
| | Reverse | 5'- GGA ACC CTG ACA CCA GGT AA -3' |
| CXCL5 | Forward | 5'- CCC TTC CTC AGT CAT AGC CG -3' |
| | Reverse | 5'- CTA TGA CTT CCA CCG TAG GGC -3' |
| CCL12 | Forward | 5'- GCT ACC ACC ATC AGT CCT CAG -3' |
| | Reverse | 5'- GAC ACT GGC TGC TTG TGA TTC -3' |
| CCL28 | Forward | 5'- CAA GCA GGG CTC ACA CTC AT -3' |
| | Reverse | 5'- GGC CAT GGG AAG TAT GGC TT -3' |
| IL17b | Forward | 5'- GGA CTG GCC GCA CAG C -3' |
| | Reverse | 5'- GCC TCC CTT GCC CTT TTC TT -3' |
| IL10 | Forward | 5'- AGG CGC TGT CAT CGA TTT CT -3' |
| | Reverse | 5'- ATG GCC TTG TAG ACA CCT TGG -3' |

**Data S2: List of Antibodies**

| Target | Isotype | Conjugate | Clone | Company | Cat # |
|---|---|---|---|---|---|
| ki67 | Rabbit IgG | Unconjugated | - | Abclonal | A16919 |
| CD4 | Rat IgG2b, κ | Alexa Fluor® 700 | GK1.5 | BioLegend | 100429 |
| CD8a | Rat IgG2a, κ | Pacific Blue™ | 53-6.7 | BioLegend | 100728 |
| CD86 | Rat IgG2a, κ | Alexa Fluor® 488 | GL-1 | BioLegend | 105017 |
| CD163 | Rat IgG2a, κ | PE | S15049F | BioLegend | 156703 |
| CD45 | Rat IgG2b, κ | Alexa Fluor® 594 | 30-F11 | BioLegend | 103144 |
| CD11b | Rat IgG2b, κ | PE | M1/70 | BioLegend | 101207 |

| | | | | | |
|---|---|---|---|---|---|
| Ly-6G/Ly-6C | Rat IgG2b, κ | FITC | RB6-8C5 | BioLegend | 108405 |
| I-A/I-E | Rat IgG2b, κ | Alexa Fluor® 647 | M5/114.15.2 | BioLegend | 107617 |
| CD11c | Armenian Hamster IgG | FITC | N418 | BioLegend | 117305 |
| FOXP3 | Mouse IgG1, κ | Alexa Fluor® 488 | 150D | BioLegend | 320011 |
| LILRB4 | Rabbit IgG | Unconjugated | - | Abclonal | A7073 |
